# Elucidating the Synergistic Role of Elm1 and Gin4 Kinases in Regulating Septin Hourglass Assembly

**DOI:** 10.1101/2023.11.08.566235

**Authors:** Joseph Marquardt, Xi Chen, Erfei Bi

## Abstract

The septin cytoskeleton is extensively regulated by post-translational modifications such as phosphorylation to achieve the diversity of architectures including rings, hourglass, and gauzes. While many of the phosphorylation events of septins have been extensively studied in the budding yeast *Saccharomyces cerevisiae*, the regulation of the kinases involved remains poorly understood. Here we show that two septin-associated kinases, the LKB1/PAR-4-related kinase Elm1 and the Nim1/PAR-1-related kinase Gin4, regulate each other at two discrete points of the cell cycle. During bud emergence, Gin4 targets Elm1 to the bud neck via direct binding and phosphorylation to control septin hourglass assembly and stability. During mitosis, Elm1 maintains Gin4 localization via direct binding and phosphorylation to enable timely remodeling of the septin hourglass into a double ring. This unique synergy ensures that septin architecture is assembled and remodeled in a temporally controlled manner to perform distinct functions during the cell cycle.

**SUMMARY:** Marquardt et al. show that the septin-associated kinases Elm1 and Gin4 regulate each other via both direct binding and phosphorylation to control septin hourglass assembly and remodeling at different points of the cell cycle in the budding yeast *Saccharomyces cerevisiae*.

## INTRODUCTION

Cells often have multiple levels of regulations to ensure fidelity from generation to generation during propagation and development, particularly in multicellular organisms. Even in the unicellular budding yeast *Saccharomyces cerevisiae*, vast gene networks cooperate to perform functions necessary to carry out its life cycle. One such pathway involves the formation of a septin collar at the neck region between a mother cell and its growing daughter/bud (Byers and Goetsch, 1976; Hartwell, 1971), which can serve as a scaffold for various cell cycle machinery and as a membrane diffusion barrier (Caudron and Barral, 2009; Gladfelter et al., 2001; Marquardt et al., 2019; McMurray and Thorner, 2009). The septins are a family of GTP-binding proteins that are conserved from yeast to humans, with the exception of land plants (Pan et al., 2007). Overexpression of septins is associated with diverse cancers, and mutations in septin genes cause neurodegenerative diseases and male infertility (Dolat et al., 2014; Hall and Russell, 2004). Septins polymerize from hetero-octamers of individual subunits (Cdc3, Cdc10, Cdc11, Cdc12, and Shs1 in budding yeast) into distinct higher order structures such as rings, collars/hourglasses, and double ring (Bertin et al., 2008; Garcia et al., 2011; Mostowy and Cossart, 2012; Oh and Bi, 2011; Ong et al., 2014). The ability to transition between these higher order structures is critical for septins to perform distinct functions throughout the cell cycle and suggest that they must be extensively regulated (Marquardt et al., 2019). The septins themselves are highly stable proteins with very little change in expression (Caviston et al., 2003; Dobbelaere et al., 2003), suggesting that any regulation occurring during the cell cycle arises from post-translational modifications of the septins themselves or septin-associated proteins (SAPs) (Marquardt et al., 2021; McMurray and Thorner, 2009).

The most common post-translational modification to regulate septins and other cellular pathways is phosphorylation by protein kinases. Indeed, nearly every architectural change of septins involves some form of phosphorylation. Prior to bud emergence, the small GTPase Cdc42 and its effectors Gic1, Gic2, and p21-activated protein kinase (PAK), Cla4, controls septin recruitment and nascent ring assembly (Caviston et al., 2003; Cvrcková et al., 1995; Gladfelter et al., 2002; Gladfelter et al., 2004; Goehring et al., 2003; Iwase et al., 2006; Sadian et al., 2013). Cla4 was subsequently found to directly phosphorylate septins, most likely Cdc10, to control the septin collar formation during bud emergence (Versele and Thorner, 2004). The LKB1/PAR-4-like kinase Elm1 has also been shown to stabilize the septin hourglass at the bud neck (Bouquin et al., 2000; Marquardt et al., 2020; Sreenivasan et al., 2003), however, this regulation may be indirect as Elm1 does not seem to phosphorylate septins directly (Versele and Thorner, 2004), but is involved in the regulation of septin architecture via the SAP Bni5 (Marquardt et al., 2020; Patasi et al., 2015). A third kinase found to regulate the stability of the septin hourglass is the Nim1/PAR-1-related kinase Gin4 that can directly bind to and phosphorylate the septin Shs1 (Asano et al., 2006; Mortensen et al., 2002), which may be important for the switch from an early hourglass to a transitional gauze-like septin structure prior to cytokinesis (Garcia et al., 2011; Ong et al., 2014). Finally, phosphorylation of Cdc3 by the cyclin-dependent kinase (CDK) Cdc28 is critical for the disassembly of septin rings after cytokinesis before a new ring is formed at the next budding site (Tang and Reed, 2002). Taken together, kinase-mediated regulations of septins and SAPs are critically important for regulating the transitions and stability of septin architecture throughout the cell cycle.

The extensive network of septin regulation by kinases becomes more complicated after examining how these kinases can regulate each other. Since Cla4, Elm1, and Gin4 show increased activity as cells pass through mitosis, it is not surprising that Cdc28 coupled to mitotic cyclins is critical for the activity of each kinase (Altman and Kellogg, 1997; Bouquin et al., 2000; Mortensen et al., 2002; Sreenivasan and Kellogg, 1999; Tjandra et al., 1998). Genetic analyses suggest that Cla4 and Elm1 both act upstream to fully activate Gin4 (Sreenivasan and Kellogg, 1999; Tjandra et al., 1998), and that Elm1 most likely acts upstream to fully activate Cla4 (Sreenivasan and Kellogg, 1999). This is also supported by the fact that *elm1τι* cells exhibit the most penetrant septin defects among the three kinases. Cla4 does not localize to the bud neck during bud growth and affects septin assembly during nascent ring formation (Gladfelter et al., 2004; Holly and Blumer, 1999; Versele and Thorner, 2004); therefore, any effect on Elm1 and Gin4 is most likely indirect because these kinases localize predominantly to the neck region in a septin-dependent manner during bud growth (Bouquin et al., 2000; Longtine et al., 1998a). There is evidence that Elm1 directly phosphorylates, activates, and properly localizes Gin4 in mitosis and that Gin4 may lead to further activation of Elm1 (Asano et al., 2006), however, the precise nature of how Elm1 and Gin4 could regulate each other and how that influences septin architecture throughout the cell cycle remains elusive.

Here we report that Gin4 and Elm1 show partially overlapping co-localization with the septins throughout bud growth. Gin4 exhibits septin co-localization prior to bud emergence and is required for the proper localization of Elm1 after bud emergence. This provides the first evidence indicating that Gin4 acts upstream of Elm1 prior to mitotic onset, primarily through direct binding and phosphorylation. As expected from a previous suggestion (Asano et al., 2006), Elm1 is critically important for maintaining Gin4 localization during later stages of mitosis and before the onset of cytokinesis. Together, these data demonstrate that the two septin hourglass-associated kinases, Gin4 and Elm1, exert mutual control to regulate hourglass assembly and remodeling during different stages of the cell cycle.

## RESULTS

### The spatiotemporal dynamics of Elm1 and Gin4 at the division site during the cell cycle

To understand if and how the kinases Elm1 and Gin4 (**Fig. 1 A**) regulate each other, we first determined their localization patterns in relation to the septins throughout the cell cycle. As seen previously (Marquardt et al., 2020), Elm1-GFP only faintly localized to the septin ring prior to bud emergence but began to accumulate during the septin hourglass stage (**Fig. 1, B and C**). This localization was maintained until 15-20 minutes prior to the septin hourglass to double ring (HDR) transition, whereby it disappeared gradually over the span of 15 minutes (**Fig. 1, D and E**). In contrast, Gin4-GFP arrived at the bud neck concurrently with the septins prior to bud emergence when a nascent septin ring was forming (**Fig. 1, B and C**). Gin4 remained at the bud neck throughout the septin hourglass stage and rapidly disassociated from the bud neck 6-8 minutes prior to the HDR transition (**Fig. 1, D and E**).

**Figure 1.**
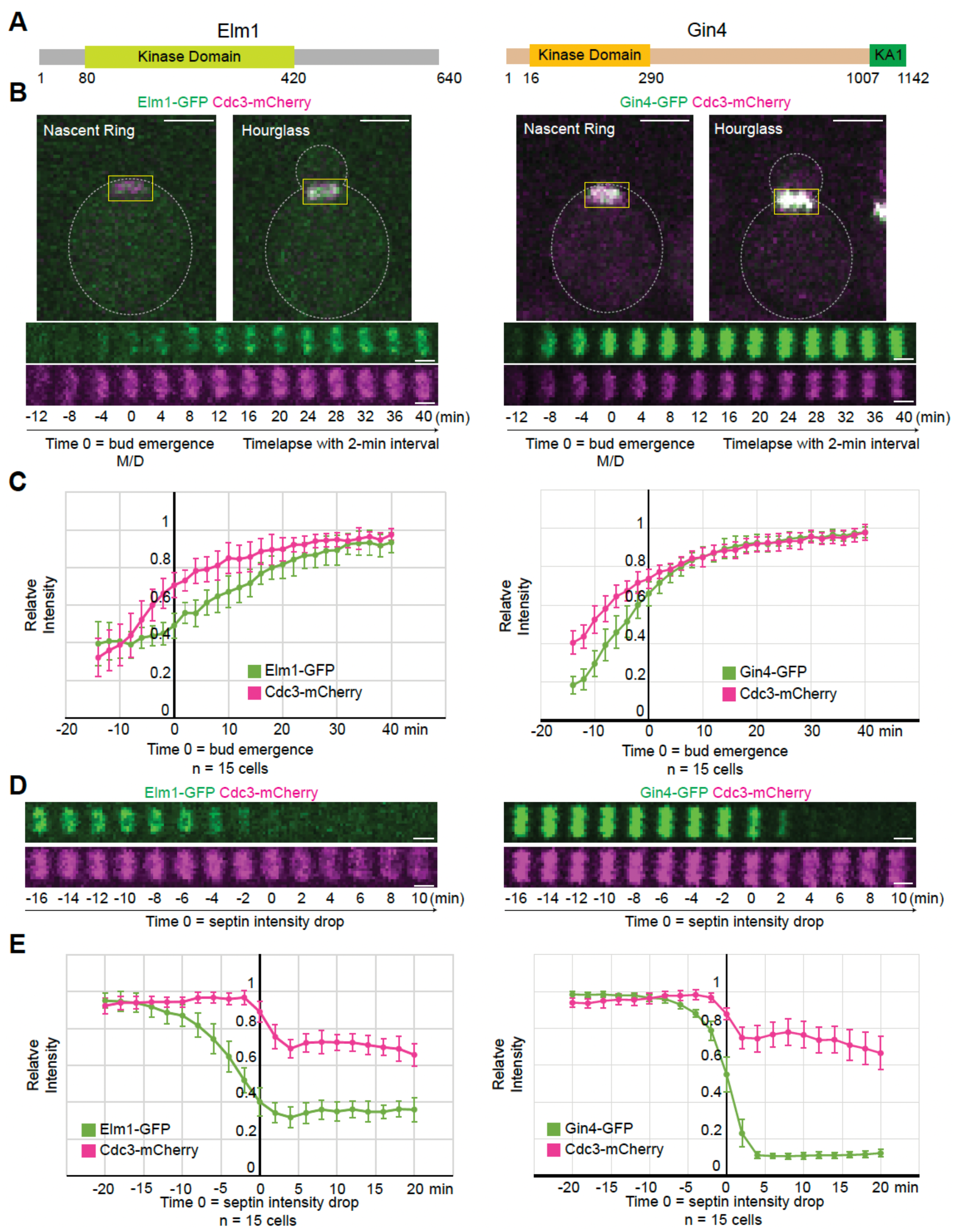
The spatiotemporal dynamics of Elm1 and Gin4 localization during the cell cycle. (A) Protein maps of Elm1 (left) and Gin4 (right) with relevant domains labeled. Numbers indicate amino acids of labeled domain boundaries. (B) (Top) Maximum-intensity projection images of representative WT cells with either Elm1-GFP (left, strain YEF9305) or Gin4-GFP (right, strain YEF10441) in green and Cdc3-mCherry (septin) in magenta at the nascent septin ring and septin hourglass stages. Gray dashed line is the cell periphery, yellow boxed region is the zoomed-in area used to create montages on the bottom, scale bar = 2 μm. (Bottom) Montages of yellow boxed region in cells shown in the top showing maximum-intensity projections of Elm1-GFP or Gin4-GFP in green and Cdc3-mCherry in magenta from 12 minutes before to 40 minutes after bud emergence with selected frames from time-lapse series taken with a 2-minute interval. For this and all subsequent montages, the mother (M) side is to the left and the daughter (D) side is to the right. T = 0 is bud emergence unless indicated otherwise. Scale bars = 1 μm. (C) Quantification of cells in Figure 1 B. Shown is background subtracted intensity of Elm1-GFP (left) or Gin4-GFP (right) in green and Cdc3-mCherry in magenta relative to the maximum value measured from the sum projection of given number cells for each strain. The mean is plotted with error bars being the standard deviation. (D) Montages of representative WT cells showing maximum-intensity projections of Elm1-GFP (left, strain YEF9305) or Gin4-GFP (right, YEF10441) in green and Cdc3-mCherry in magenta from 16 minutes before to 10 minutes after septin HDR from time-lapse series taken with a 2-minute interval. T = 0 is septin HDR as indicated by septin intensity drop. Scale bars = 1 μm. (E) Quantification of cells in Figure 1 D. Shown is background subtracted intensity of Elm1-GFP (left) or Gin4-GFP (right) in green and Cdc3-mCherry in magenta relative to the maximum value measured from the sum projection of given number cells for each strain. The mean is plotted with error bars being the standard deviation.

We confirmed this temporal difference in the same cell by using a strain expressing Elm1-GFP and Gin4-mScarlet. Gin4 arrived earlier than Elm1 prior to bud emergence (**Fig. S1, A and B**) and was maintained at the bud neck approximately 6-8 minutes longer than Elm1 just prior to cytokinesis (as indicated by mitotic spindle break) (**Fig. S1, C and D**). The above data provide temporal evidence that Gin4 arrives prior to Elm1 and is in prime position to act upstream of Elm1 prior to the septin hourglass stage.

### Gin4 is required for localization but not full function of Elm1 during the septin hourglass stage

To determine the localization dependency and functional relationship between Elm1 and Gin4 during the cell cycle, we generated a precise deletion for each kinase and examined its impact on both the septins and the alternate kinase. As expected (Bouquin et al., 2000; Marquardt et al., 2020), nearly all of the *elm1τι* cells were highly elongated with septins beginning to migrate to the growing bud tip 15-20 minutes after bud emergence (**Fig. 2 A**), suggesting a critical role of Elm1 in controlling septin hourglass stability. Gin4-GFP showed no change in its initial recruitment to the presumed bud site in *elm1τι* cells but started to migrate with the septins into the bud tip ∼15 minutes after bud emergence (**Fig. 2, A and B**). Thus, it appears that Elm1 does not change the ability of Gin4 to localize and thereby interact with the septins at this stage of the cell cycle. Gin4 has been reported to be deficient in interacting with the septin Cdc11 in the absence of Elm1 or the septin Shs1 (Mortensen et al., 2002). To compare this effect in live cells, we investigated Gin4-GFP in *shs1Δ* cells. While the kinetic signature of Gin4 recruitment remained unchanged in comparison to wild-type (WT) cells, there was a 30% drop in total intensity measured during this time, which was not seen in *elm1Δ* cells (**Fig. 2, C-E**). This data indicates that our observation of the normal recruitment of Gin4 to septin structures during the early stage of budding in *elm1Δ* cells is bona fide.

**Figure 2.**
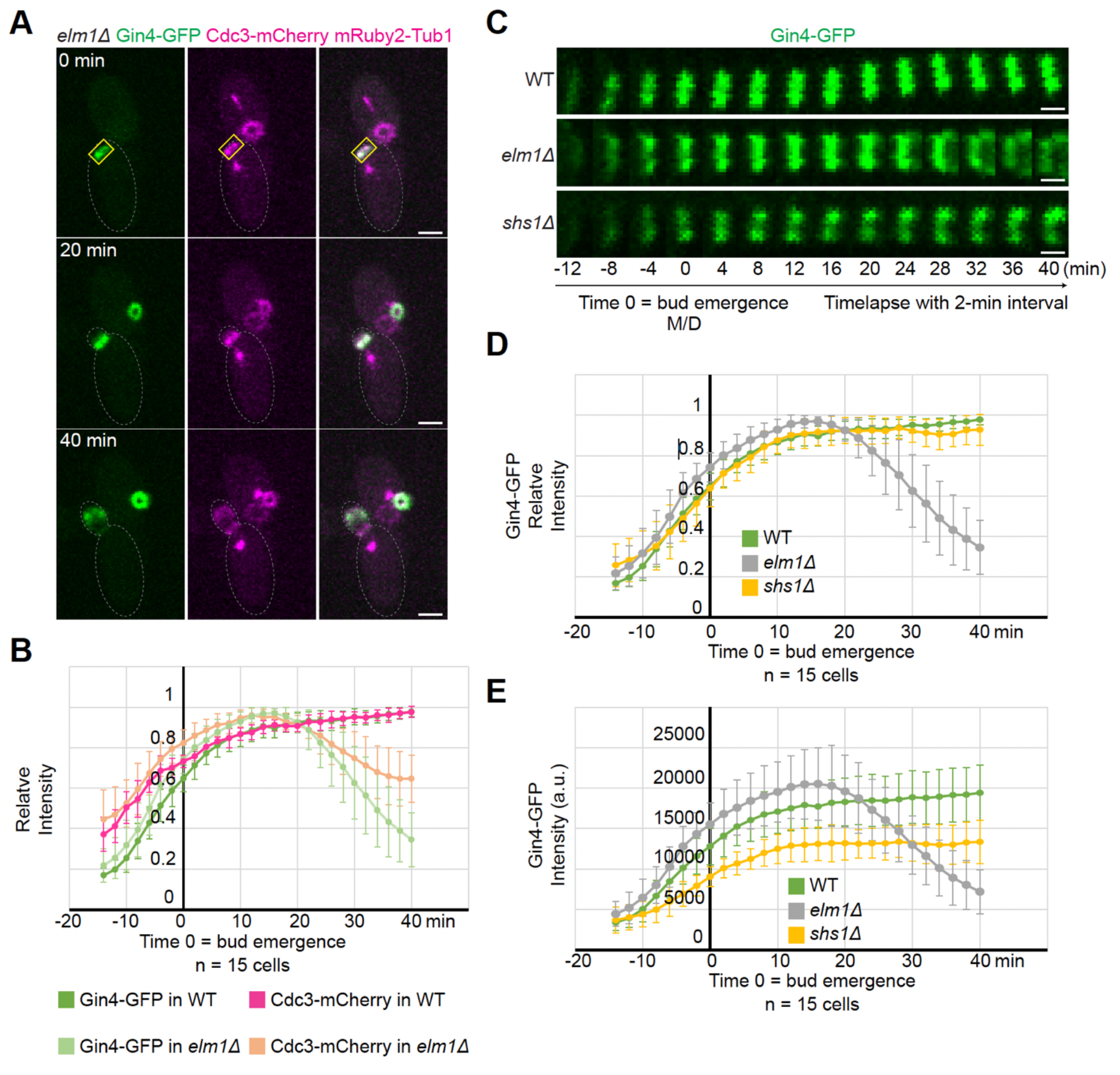
Gin4 is displaced from the bud neck concurrently with septins in *elm1Δ* cells. (A) Maximum-intensity projection images of representative YEF10559 cells (*elm1Δ GIN4-GFP CDC3-mCherry mRuby2-Tub1*) from a time-lapse series taken with a 2-minute interval with Gin4-GFP in green (left), Cdc3-mCherry and mRuby2-Tub1 (tubulin) in magenta (center), and merged (right) at indicated times. T = 0 is bud emergence, gray dashed line is the cell periphery, yellow boxed region is the zoomed-in area used to create montages for Figure 2 C, scale bars = 2 μm. (B) Quantification of cells in Figure 2 A and YEF10558 (*GIN4-GFP CDC3-mCherry mRuby2-Tub1*). Shown is background subtracted intensity of Gin4-GFP in dark green (WT) or light green (*elm1Δ*) and Cdc3-mCherry in magenta (WT) or tan (*elm1Δ*) relative to the maximum value measured from the sum projection of given number cells for each strain. The mean is plotted with error bars being the standard deviation. (C) Montages of Gin4-GFP in representative WT (YEF10558), *elm1Δ* (YEF10559), and *shs1Δ* (YEF11454) cells showing maximum-intensity projections from 12 minutes before to 40 minutes after bud emergence from time-lapse series taken with a 2-minute interval. T = 0 is bud emergence, scale bars = 1 μm. (D) Quantification of cells in Figure 2 C. Shown is background subtracted intensity of Gin4-GFP in WT (green), *elm1Δ* (gray), and *shs1Δ* (yellow) relative to the maximum value measured from the sum projection of given number cells for each strain. WT and *elm1Δ* curves are same values used for Figure 2 B. The mean is plotted with error bars being the standard deviation. (E) Quantification of cells in Figure 2 C. Shown is integrated measured background subtracted intensity of Gin4-GFP in WT (green), *elm1Δ* (gray), and *shs1Δ* (yellow) from the sum projection of given number cells for each strain. The mean is plotted with error bars being the standard deviation. A.U. = arbitrary units.

Only 40-50% of the *gin4Δ* cells exhibited an elongated morphology (**Fig. 3 A**). Regardless of the cell shape, the septins behaved similarly as in *elm1Δ* cells, but to a much lower degree. The septins began to dissociate from the bud neck 15-20 minutes after bud emergence, but only lost approximately 15-20% of the signal before returning to the bud neck 20 minutes later (**Fig. 3, B and C**), suggesting a role of Gin4 in controlling septin hourglass stability. This may explain why the cells show a lesser penetrance and degree of the elongated bud phenotype than in *elm1Δ* cells. In stark contrast, Elm1-GFP was largely absent from the bud neck in *gin4Δ* cells, which appeared to be the same in round or elongated cells (**Fig. 3 D**). This localization deficiency was unlikely caused by a change in protein level, as the total amount of Elm1 was reduced only by about 40% in *gin4Δ* cells (**Fig. 3 E**). The localization dependency data supports the notion that Gin4 acts upstream of Elm1 at the beginning of the cell cycle. Importantly, these observations also suggest that the neck localization alone cannot explain the full function of Elm1 in controlling septin hourglass stability during the cell cycle.

**Figure 3.**
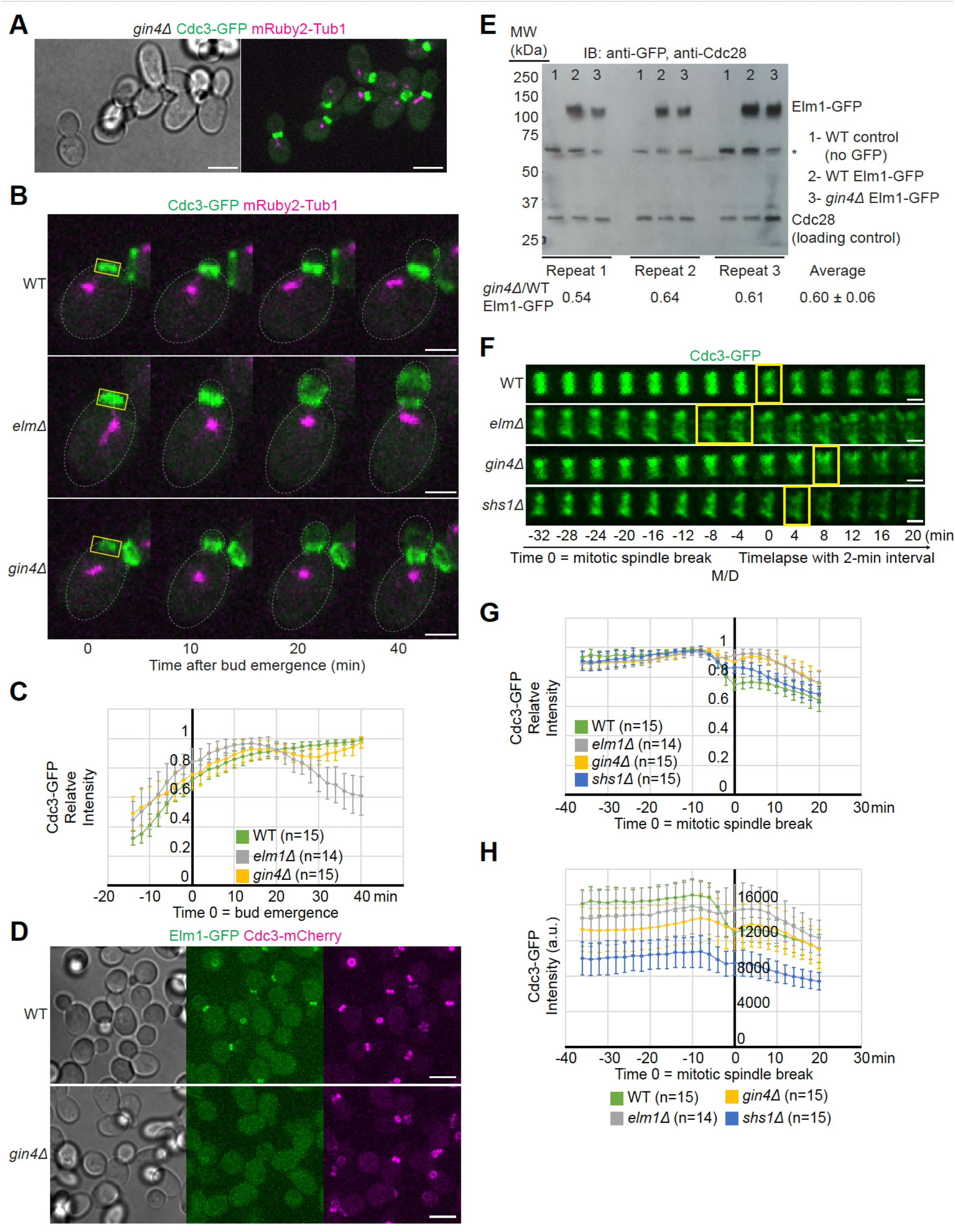
Septins exhibit mild bud neck displacement while Elm1 is absent from the bud neck in *gin4Δ* cells. (A) Representative images of YEF9641 (*gin4Δ CDC3-GFP mRuby2-Tub1*) cells with brightfield (left) and maximum intensity projection of merged Cdc3-GFP in green and mRuby2-Tub1 in magenta (right). Scale bar = 5 μm. (B) Maximum-intensity projection images of representative WT (YEF9180), *elm1Δ* (YEF9935), and *gin4Δ* (YEF9641) cells from a time-lapse series taken with a 2-minute interval with Cdc3-GFP in green and mRuby2-Tub1 in magenta at indicated times. T = 0 is bud emergence, gray dashed line is the cell periphery, yellow boxed region is the area used for measurements in Figure 3 C, scale bars = 2 μm. (C) Quantification of cells in Figure 3 B. Shown is background subtracted intensity of Cdc3-GFP in WT (green), *elm1Δ* (gray), and *gin4Δ* (yellow) relative to the maximum value measured from the sum projection of given number cells for each strain. The mean is plotted with error bars being the standard deviation. (D) Representative images of Elm1-GFP and Cdc3-mCherry in WT (YEF10440) and *gin4Δ* (YEF10460) cells with brightfield (left) and maximum intensity projection of Elm1-GFP in green (middle) and Cdc3-mCherry in magenta (right). Scale bars = 5 μm. (E) (Top) Immunoblot (IB) analysis of Elm1-GFP and Cdc28 from three independent replicates of total protein isolates from WT (YEF9327, sample 1), Elm1-GFP (YEF10749, sample 2), and *gin4Δ* Elm1-GFP (YEF10750, sample 3) cells. MW = protein marker with indicated sizes in kilo Daltons (kDa) at indicated positions. Asterisk (*) indicates a non-specific band from the anti-GFP antibody. (Bottom) Quantification of ratio of Elm-GFP/Cdc28 band intensity in *gin4Δ* samples relative to that of WT Elm1-GFP samples for each replicate. Average is presented from the three replicates ± standard deviation. (F) Montages of representative WT (YEF9180), *elm1Δ* (YEF9935), *gin4Δ* (YEF9641), and *shs1Δ* (YEF8438) cells showing maximum-intensity projections of Cdc3-GFP in green from 32 minutes before to 20 minutes after mitotic spindle break with selected frames from time-lapse series taken with a 2-minute interval. T = 0 is mitotic spindle break. Scale bars = 1 μm. (G) Quantification of cells in Figure 3 F. Shown is background subtracted intensity of Cdc3-GFP in WT (green), *elm1Δ* (gray), *gin4Δ* (yellow), and *shs1Δ* (blue) relative to the maximum value measured from the sum projection of given number cells for each strain. The mean is plotted with error bars being the standard deviation. (H) Quantification of cells in Figure 3 F. Shown is background subtracted intensity of Cdc3-GFP in WT (green), *elm1Δ* (gray), *gin4Δ* (yellow), and *shs1Δ* (blue) relative to the region of interest’s area measured from the sum projection of given number cells for each strain. The mean is plotted with error bars being the standard deviation. A.U. = arbitrary units.

Gin4 had previously been described to have a function in the HDR transition (Asano et al., 2006), but a detailed investigation of its regulatory role during this transition has not been investigated. Time-lapse imaging of *gin4Δ* cells revealed that the septins underwent the HDR transition slightly later than in WT cells (**Fig. 3F**, yellow boxes). The septins also failed to disassemble and lose 30% of their intensity during the transition that has been seen previously (Dobbelaere et al., 2003; Wloka et al., 2011) and instead only lost 10-15% of their maximum intensity (**Fig. 3, F and G**). This reduction in signal loss was similar to our previous results using *elm1Δ* and *shs1Δ* cells (Marquardt et al., 2020) and a direct comparison showed that Cdc3-GFP raw intensity in *elm1Δ* and *gin4Δ* cells was 8% and 16% lower than that of WT cells 10 minutes prior to mitotic spindle breakage, respectively (**Fig. 3 H**). This indicates that Gin4 regulates the timely reorganization of the septin hourglass structure to double ring during cytokinesis.

### Gin4 controls Elm1 localization via direct interaction and phosphorylation

Since Gin4 could affect Elm1 localization by directly binding to or phosphorylating Elm1, we investigated both possibilities. Firstly, we examined the localization of Elm1 in cells expressing Gin4^ΔKA1^, which lacks the C-terminal KA1 domain known to be required for membrane binding and bud neck localization (Moravcevic et al., 2010), from the endogenous *GIN4* locus. Not surprisingly, Elm1 was completely absent from the bud neck in these cells (**Fig. 4 A**), suggesting that the membrane binding and/or neck localization of Gin4 is required for the association of Elm1 with the septin hourglass. The simplest possibility for the observation is that Gin4 recruits Elm1 to the bud neck via a direct interaction. Indeed, we found that GST-Elm1 was able to interact with 6xHis-SUMO-Gin4 in vitro (**Fig. 4 B**). Strikingly, the C-terminal non-kinase domain of Elm1 displayed strong interaction with Gin4 (**Fig. 4 B**, bottom lane 4), whereas its kinase domain did not exhibit any binding capacity (**Fig. 4 B**, bottom lane 3). This is consistent with the previous observation that the non-kinase domain of Elm1 is responsible for its bud neck localization (Marquardt et al., 2020; Moore et al., 2010). Thus, we have established a direct interaction between Gin4 and Elm1.

**Figure 4.**
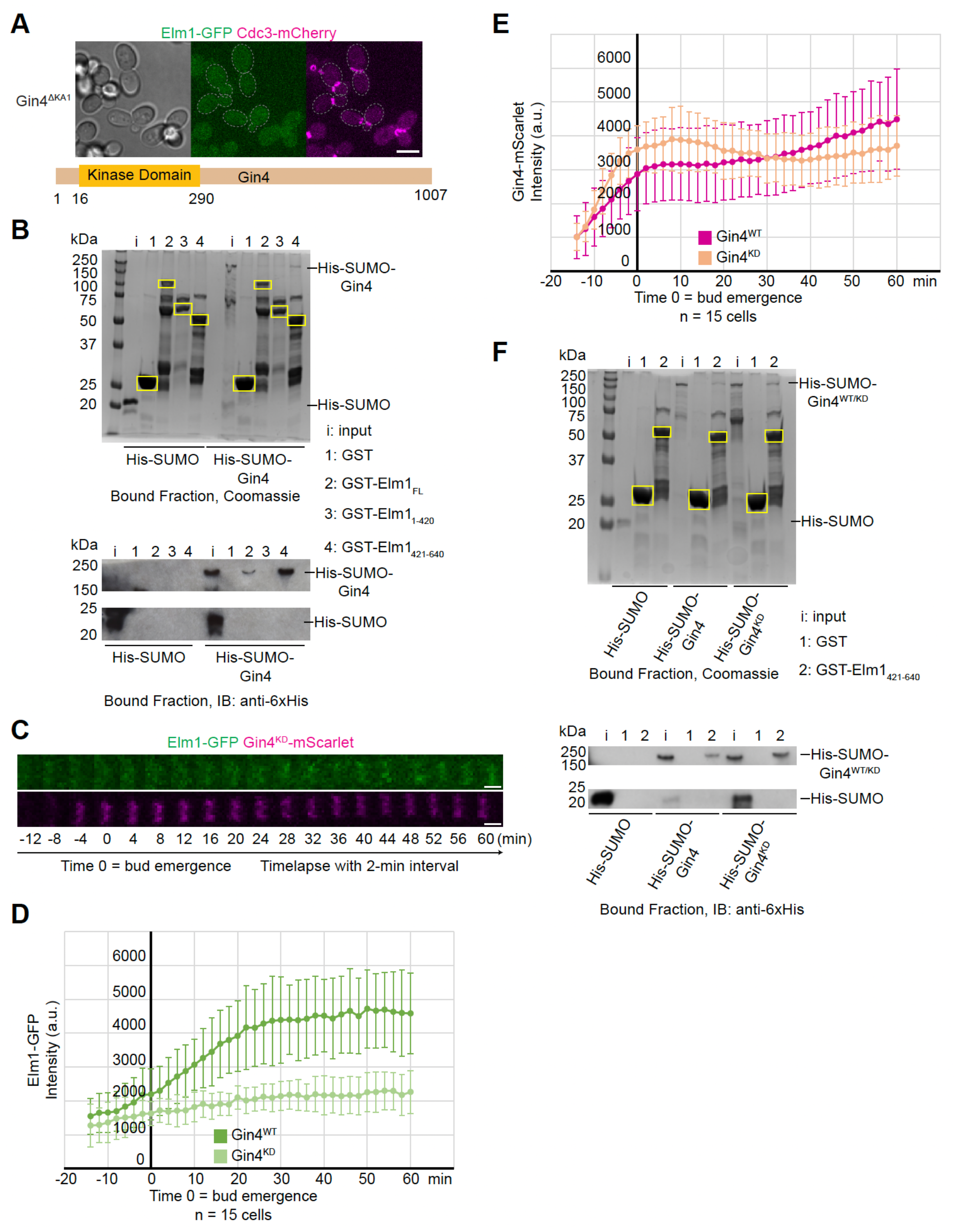
Gin4 regulates Elm1 localization through direct binding and phosphorylation. (A) Representative image of YEF10672 (*gin4^ΔKA1^ ELM1-GFP CDC3-mCherry*) cells with brightfield (left) and maximum intensity projection of Elm1-GFP in green (middle) and Cdc3-mCherry in magenta (right). Gray dashed line is cell periphery. Scale bar = 5 μm. (B) (Top) In vitro binding assay results for indicated GST-tagged proteins bound to glutathione resin and indicated 6xHis-SUMO-tagged protein separated by SDS-PAGE and Coomassie Blue stained. Yellow boxes indicate purified GST-tagged protein. (Bottom) In vitro binding assay results for indicated GST-tagged proteins bound to glutathione resin and indicated 6xHis-SUMO-tagged protein separated by SDS-PAGE and immunoblotted with anti-6xHis antibody. kDa = kiloDalton. This experiment was repeated 3 times with consistent interaction detected for the full-length Elm1, strong interaction for its C-terminal fragment, and weak or no interaction for its N-terminal fragment. (C) Montages of representative YEF10673 (Elm1-GFP in green and Gin4^KD^-mScarlet in magenta) cells showing maximum-intensity projections from 12 minutes before to 60 minutes after bud emergence from time-lapse series taken with a 2-minute interval. T = 0 is bud emergence, scale bars = 1 μm. (D) Quantification of Elm1-GFP signal from cells in Figure 4 C. Shown is integrated measured background subtracted intensity in Gin4^WT^ (YEF10802, green) and Gin4^KD^ (light green) from the sum projection of given number cells for each strain. The mean is plotted with error bars being the standard deviation. A.U. = arbitrary units. (E) Quantification of Gin4-mScarlet signal from cells in Figure 4 C. Shown is integrated measured background subtracted intensity in Gin4^WT^ (YEF10802, magenta) and Gin4^KD^ (tan) from the sum projection of given number cells for each strain. The mean is plotted with error bars being the standard deviation. A.U. = arbitrary units. (F) (Top) In vitro binding assay results for indicated GST-tagged proteins bound to glutathione resin and indicated 6xHis-SUMO-tagged protein separated by SDS-PAGE and Coomassie Blue stained. Yellow boxes indicate purified GST-tagged protein. (Bottom) In vitro binding assay results for indicated GST-tagged proteins bound to glutathione resin and indicated 6xHis-SUMO-tagged protein separated by SDS-PAGE and immunoblotted (IB) with anti-6xHis antibody. kDa = kiloDalton. This experiment was repeated 2 times with indistinguishable interaction differences detected between 6xHis-SUMO-Gin4WT and 6xHis-SUMO-Gin4KD to the C-terminal non-kinase domain of Elm1.

Next, we examined the possibility of Gin4 regulating Elm1 localization through phosphorylation. In cells harboring a mutation at the endogenous locus known to largely abolish the kinase activity of Gin4 (i.e., the *gin4-K48A* allele with its protein product known as Gin4^KD^), Elm1-GFP showed a significantly weakened and delayed signal that flashed between time points, suggesting an instability for the bud neck-localized Elm1 (**Fig. 4, C and D**). This was observed despite the fact that Gin4^KD^ exhibited largely the same intensity at the bud neck throughout the septin hourglass stage (**Fig. 4, C and E**), and that Gin4^KD^ did not show any decrease in its direct binding to the non-kinase domain of Elm1 than the WT Gin4 (hereafter called Gin4^WT^) in vitro (**Fig. 4 F**). These observations suggest that the direct binding of Gin4 to Elm1 may be necessary but not sufficient for Elm1 localization, which raises the possibility that Gin4 may also phosphorylate Elm1 to enable its localization. To test this possibility, we immunoprecipitated Elm1-GFP from both WT and *gin4Δ* cells and examined the phosphorylation status of Elm1 via LC-MS/MS analysis (**Fig. S2 A**). Six total sites in Elm1 were found to be differentially phosphorylated between the two sample sets (**Fig. S2 B**). The five sites found enriched in the WT sample (S30, S96, S152, S519, and T551) constitute the possible in vivo set of phosphorylation sites by Gin4. However, Gin4 may not directly phosphorylate these sites, e.g., Gin4 could regulate the ability of other kinases to access Elm1 in vivo. Therefore, we performed an in vitro kinase reaction with purified 6xHis-SUMO-Gin4 and GST-Elm1^KD^ and performed the same LC-MS/MS analysis (**Fig. 5 A**). In this in vitro dataset, seven phosphorylation sites in Elm1 (S324, S422, S424, T463, S519, S604, and S605) were found only in the sample in which Gin4 was added to the kinase reaction (**Fig. 5 B**), thus, these sites represent potential direct Gin4-dependent sites. Strikingly, all but one (S324) were found to be in the bud neck localization domain and S519 was shared with our in vivo data set. During the in vitro kinase assay analysis, we also discovered a total of 63 phosphorylation sites in the sequence of 6xHis-SUMO-Gin4, which may constitute auto-phosphorylation sites (**Fig. S2 C**).

**Figure 5.**
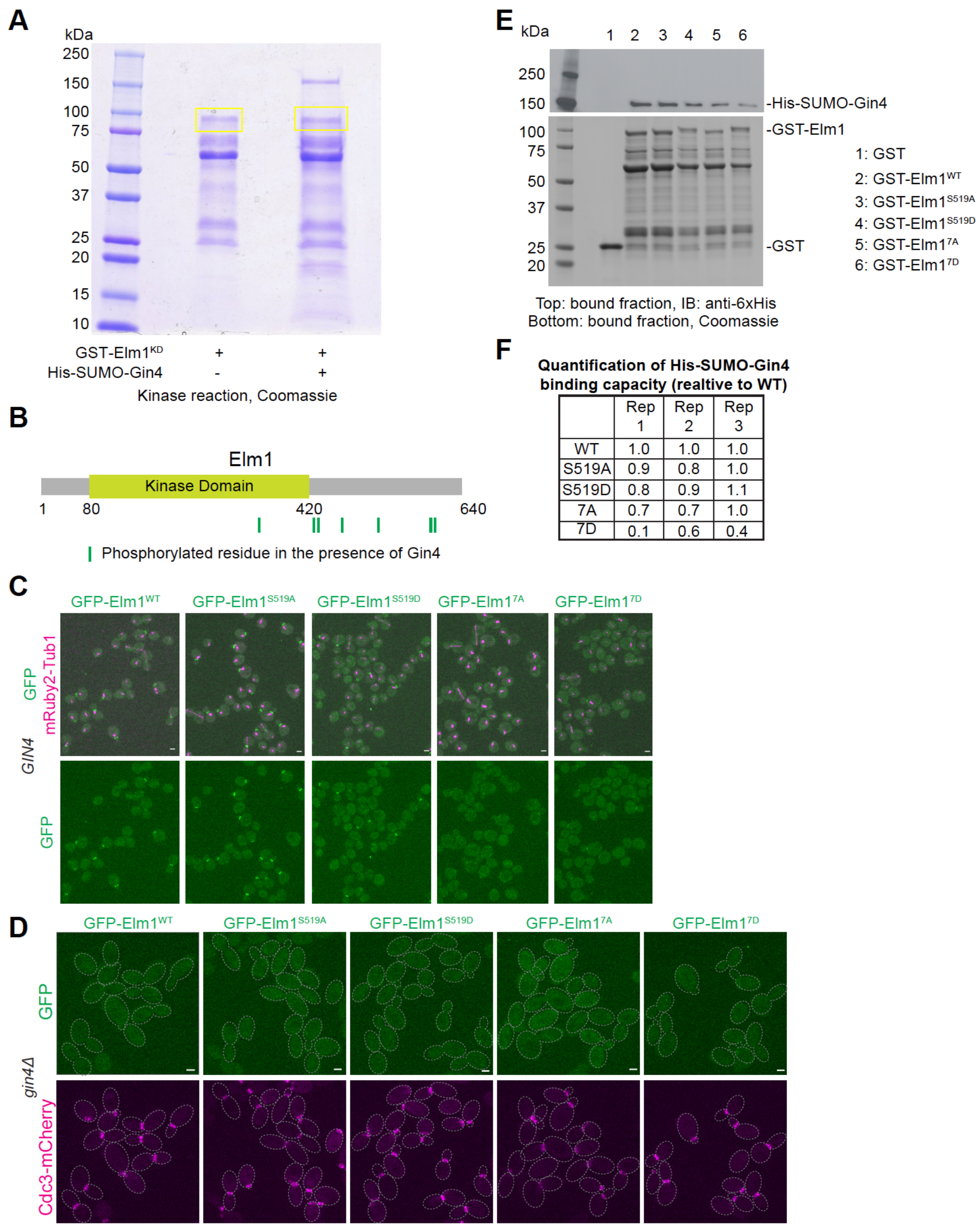
Gin4 directly phosphorylates Elm1 to regulate its bud neck localization. (A) In vitro kinase assay results for GST-Elm1^KD^ incubated with 6xHis-SUMO-Gin4 separated by SDS-PAGE and Coomassie Blue stained. Yellow boxes indicate regions excised for mass spectrometry analysis. kDa = kiloDalton. (B) Protein schematic of Elm1 with indicated domain boundaries labeled with amino acid positions. Green vertical lines indicate the position of a phosphorylated residue discovered via mass spectrometry. Those sites from left to right are S324, S422, S424, T463, S519, S604, and S605. (C) Representative images of cells for GFP-Elm1^WT^ (YEF11679), GFP-Elm1^S519A^ (YEF11680), GFP-Elm1^S519D^ (YEF11681), GFP-Elm1^7A^ (YEF11682), and GFP-Elm1^7D^ (YEF11683) in green and mRuby2-Tub1 in magenta. The images show maximum projections. Scale bars = 2 μm. (D) Representative images of the indicated *gin4Δ* cells with GFP-tagged Elm1 mutants in green and mRuby2-TUB1 in magenta. Strains used are as follows from left to right: YEF11708 (*gin4Δ GFP-ELM1^WT^*), YEF11709 (*gin4Δ GFP-elm1^S519A^*), YEF11710 (*gin4Δ GFP-elm1^S519D^*), YEF11711 (*gin4Δ GFP-elm1^7A^*), and YEF11712 (*gin4Δ GFP-elm1^7D^*). The images show maximum projections. The dotted line indicates the cell periphery. Scale bars = 2 µm. (E) In vitro binding assay results for the indicated GST-tagged proteins bound to glutathione resin and their ability to pull down His-SUMO-Gin4. Top: immunoblotted with antibody against 6x-His; Bottom: Coomassie Blue-stained. kDa = kiloDalton. This experiment was repeated 3 times and a representative immunoblot is shown. (F) Quantification of GST-Elm1 mutants binding ability to 6xHis-SUMO-Gin4. The formula used to calculate values = (membrane band intensity of Gin4/gel band intensity of Elm1)/WT calculated value.

Since S519 was found in both the in vivo and in vitro potential phosphorylation datasets, we began testing the biological significance of phosphorylation at this site. Mutating this serine residue to either alanine or aspartic acid at the genomic locus to mimic the dephosphorylated or phosphorylated condition, respectively, had no significant effect on the localization timing or amount during the septin hourglass stage (**Fig. 5 C and Fig. S3, A and B**). During the septin HDR transition, both mutants (Elm1^S519A^ and Elm1^S519D^) exhibited a mild, yet significant decrease in their overall localization capacity to the late hourglass structure (**Fig. S3, C and D**). To examine if Elm1^S519D^ could bypass the requirement for Gin4 phosphorylation, we tested its localization in *gin4Δ* cells. Surprisingly, it behaved identically as both Elm1^WT^ and Elm1^S519A^, with almost no localization at the bud neck region (**Fig. 5 D**). This suggests that either multiple Gin4 phosphorylation sites are required for biological significance or that the regulation is a combination of both phosphorylation and direct binding to Gin4.

To address these possibilities, we mutated all residues found in the in vitro dataset to either alanine or aspartic acid at the genomic locus. Since these sites represented the most likely direct Gin4-mediated phosphorylation sites (unlike the residues uncovered in the in vivo dataset), we only focused on these as genuine phosphorylation candidates. As expected, the Elm1^7A^ mutant exhibited a near complete loss of bud neck localization in otherwise WT or *gin4Δ* cells (**Fig. 5, C and D**). Surprisingly, the Elm1^7D^ mutant exhibited a drastic decrease in bud neck localization in WT cells and a complete loss in bud neck localization in *gin4Δ* cells (**Fig. 5, C and D**). There was no significant change in total expression of any mutant GFP-Elm1 protein to explain this loss of bud neck localization (**Fig. S3 E**). To address why Elm1^7D^, which should not require Gin4-mediated phosphorylation to be recruited to the bud neck, would exhibit the same phenotype as the alanine substitution mutant, we examined the in vitro binding capacity of all our mutant Elm1 constructs with Gin4. Only Elm1^7D^ showed a reliable decrease in binding capacity to Gin4 (**Fig. 5, E and F**). This explains why this mutant fails to localize: it has lost the ability to be targeted to the bud neck by direct binding to Gin4.

With the localization of both Elm1^7A^ and Elm1^7D^ being vastly altered when compared to that of Elm1^WT^, it was surprising to not witness any robust septin defects in these mutants. Mild septin phenotypes in various Elm1 mutants can be exacerbated by deletions of either the septin *SHS1* or the septin bundling protein *BNI5* (Marquardt et al., 2020). We therefore combined each of these deletions with all the constructed Elm1 phospho-mutants. The added deletion of *BNI5* (*bni5Δ*) had no effect on the growth of cells (**Fig. S4 A**) or the septins during the cell cycle at either 25°C or the elevated 37°C (**Fig. S4, B and C**). This is a significant finding that allows us to conclude that all our constructed mutants have retained Elm1 kinase activity since it has already been shown that *bni5Δ* exhibits a strong septin and cell morphology phenotype in the absence of Elm1 activity (Marquardt et al., 2020). Additionally, Elm1^7D^ failed to localize to the bud neck in *bni5Δ* cells (**Fig. S4 B**). Elm1 and Bni5 are known to directly interact with one another, but Elm1 localization is not affected by a deletion of *BNI5* (Marquardt et al., 2020; Patasi et al., 2015). This implies that the faint localization of Elm1^7D^ in otherwise WT cells, which showed reduced binding to Gin4, may require Bni5 for bud neck localization. While *shs1Δ* in combination with all constructed Elm1 mutants displayed no obvious growth defects (**Fig. 6 A**), the *elm1^7A^*mutant allele combination exhibited a very drastic cell elongation phenotype that was coupled with septin abnormalities (**Fig. 6 B**). Specific septin phenotypes included asymmetric localization to the daughter side of the bud neck and abnormal localization at the bud cortex in 20% and 36.7% of cells, respectively (**Fig. 6, C and D**). The finding that Elm1^7D^ resulted in slightly elongated cell morphology and less penetrant septin phenotypes in *shs1Δ* cells (**Fig. 6, B - D**) provides evidence that the Gin4-mediated phosphorylation sites have a biological significance when other septin perturbations are present, but these phosphorylation events must be dynamic for proper septin regulation.

**Figure 6.**
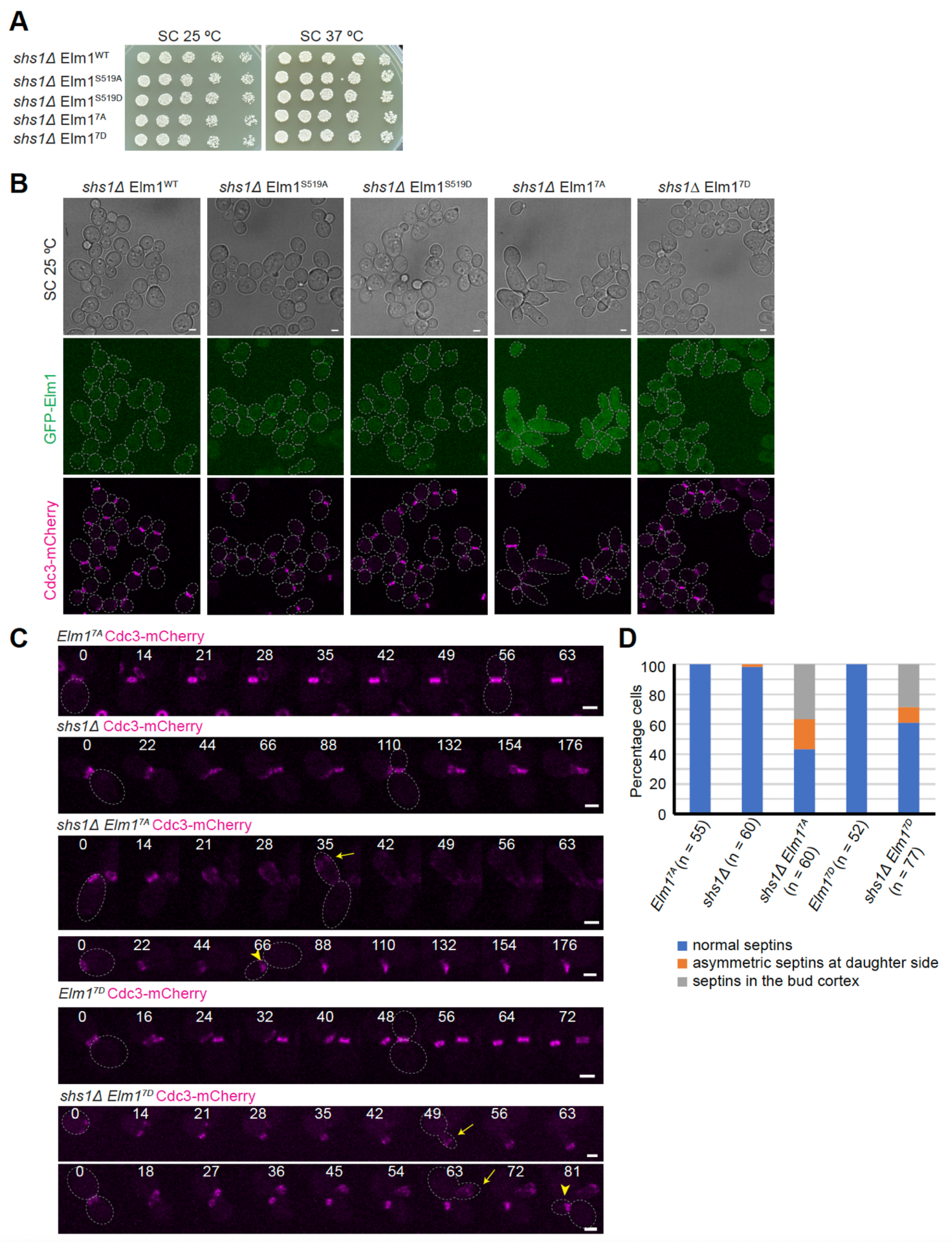
Gin4-mediated phosphorylation of Elm1 is required for normal septin morphology in the absence of *SHS1*. (A) The sensitivity of *shs1Δ* cells with Elm1 mutants was tested on synthetic complete (SC) plates. Ten-fold serial dilutions of YEF11766 (*shs1Δ GFP-ELM1^WT^ CDC3-mCherry*), YEF11767 (*shs1Δ GFP-elm1^S519A^ CDC3-mCherry*), YEF11768 (*shs1Δ GFP-elm1^S519D^ CDC3-mCherry*), YEF11769 (*shs1Δ GFP-elm1^7A^ CDC3-mCherry*), and YEF11770 (*shs1Δ GFP-elm1^7D^ CDC3-mCherry*) cultures were spotted on SC plates and incubated at 25°C (left) and 37°C (right) for 3 days. (B) Representative images of the indicated *shs1Δ* cells with brightfield (top) and maximum intensity projections of GFP-tagged Elm1 mutants shown in green (middle) and Cdc3-mCherry shown in magenta (bottom). Strains used are listed in Figure 6 A. The images show maximum projections. The gray dashed line indicates the cell periphery. Scale bars = 2 µm. (C) Montages of representative cells of the indicated strains, with CdC3-mCherry shown in magenta. Strains used are as follows from top to bottom: YEF11691 (*GFP-elm1^7A^ CDC3-mCherry)*, YEF11766 (*shs1Δ GFP-ELM1^WT^ CDC3-mCherry*), and YEF11769 (*shs1Δ GFP-elm1^7A^ CDC3-mCherry*), YEF11692 (*GFP-elm1^7D^ CDC3-mCherry*), and YEF11770 (*shs1Δ GFP-elm1^7D^ CDC3-mCherry*). The images show maximum projections. The yellow arrow indicates septins localized in the bud cortex, the yellow arrowhead indicates asymmetric septins at daughter side. The gray dashed line indicates the cell periphery. Scale bars = 2 µm. (D) Quantification of cells with the indicated septin phenotypes in Figure 6 C. n = number of cells analyzed for each strain.

The above data lead us to conclude that Elm1 localization to the septin hourglass is mediated through direct binding of Gin4 to the C-terminal non-kinase domain of Elm1 and subsequent maintenance at the bud neck through direct phosphorylation by Gin4. These phosphorylation sites have biological relevance when the septins are perturbed in mutants such as *shs1Δ* cells. Collectively, this illustrates a complex regulatory system to ensure septin homeostasis.

### Artificial tethering of Elm1 to the septins can largely suppress defects in *gin4Δ* cells

To examine if a primary reason for the cell morphology and septin phenotypes of *gin4Δ* cells is the loss of Elm1 localization at the bud neck, we utilized the GFP/GBP strategy to independently target Elm1 to the septins (Kubala et al., 2010; Marquardt et al., 2020; Rothbauer et al., 2006). Remarkably, when Elm1^WT^-mApple-GBP was tethered to Shs1-GFP in *gin4Δ* cells, 82.9 ± 3.8% cells exhibited a normal septin structure with a round morphology (compared to only 48.3 ± 4.2% in untethered *gin4Δ* cells) (**Fig. 7, A and B**). When the cells were grown at 37°, the artificial targeting of Elm1^WT^-mApple-GBP rescue efficiency was lower (67.0 ± 5.2% round cells), yet still significantly improved from untethered *gin4Δ* cells (38.8 ± 5.0% round cells) (**Fig. 7, A and B**). Elm1^KD^-mApple-GBP tethered to Shs1-GFP can efficiently rescue phenotypes associated with Elm1^KD^ (Marquardt et al., 2020). While still providing a moderate rescue over untethered *gin4Δ* cells, Elm1^KD^-mApple-GBP could not rescue to the same capacity of the WT allele (**Fig. 7, C and D**). These data suggest that Gin4 regulates septin hourglass assembly and cell morphology at least in part by controlling Elm1 localization, and that both localization and kinase activity of Elm1 are required for effective suppression of *gin4Δ* cells.

**Figure 7.**
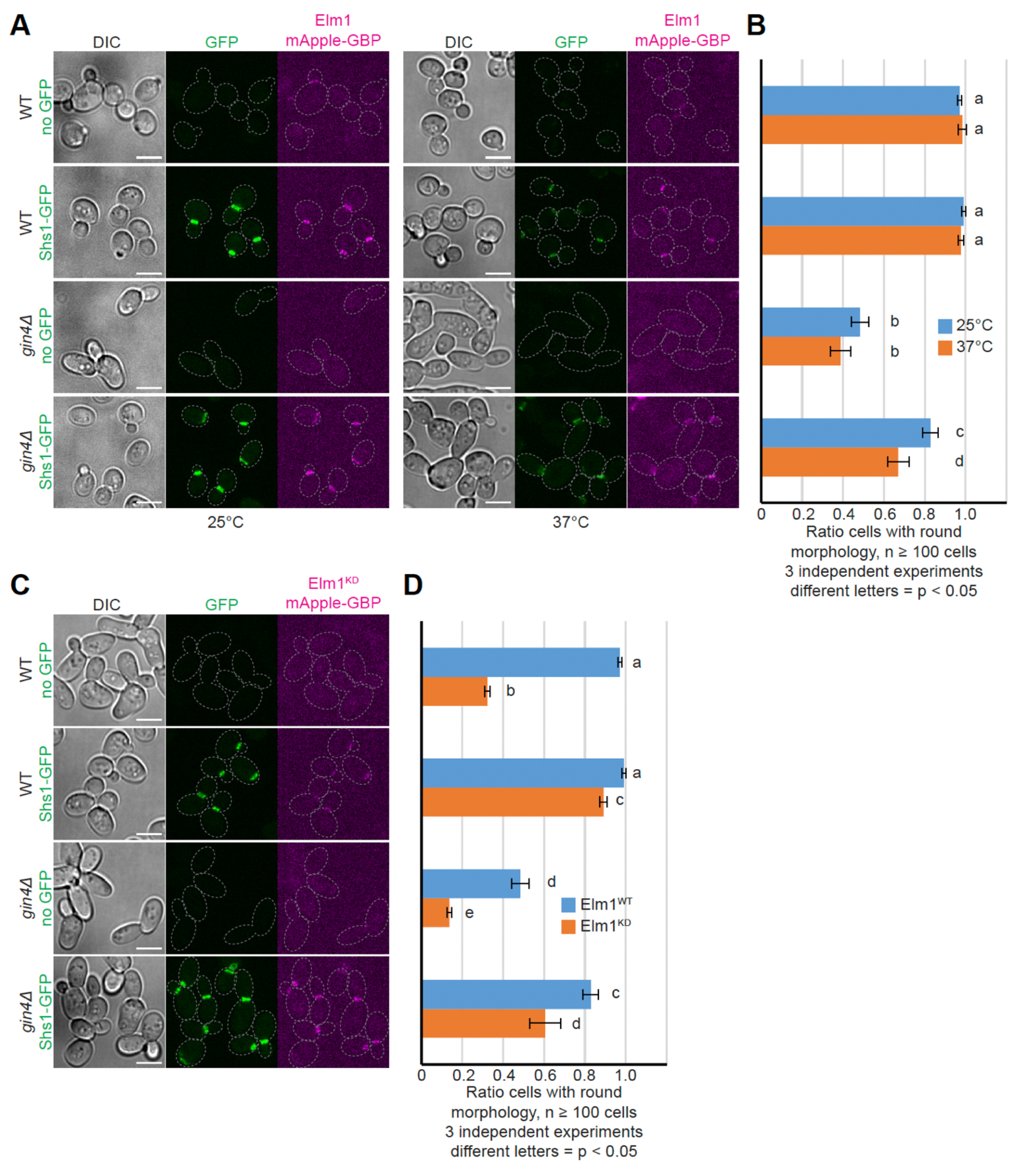
Artificial tethering of Elm1 to bud neck can rescue *gin4Δ* phenotypes. (A) Representative images of indicated cells grown overnight at either 25°C (left) or 37°C (right) with brightfield and maximum intensity projections of Shs1-GFP in green and Elm1-mApple-GBP in magenta. Strains used are as follows from top to bottom: YEF9448 (*ELM1-mApple-GBP*), YEF10665 (*SHS1-GFP ELM1-mApple-GBP*), YEF9485 (*gin4Δ ELM1-mApple-GBP*) and YEF10666 (*gin4Δ SHS1-GFP ELM1-mApple-GBP*). Gray dashed line indicates the cell periphery. Scale bars = 5 μm. (B) Quantification of the ratio of round cells in strains used in Figure 7 A. Plotted is the average of 3 independent experiments of n ≥ 100 cells. Error bars are standard deviation. Lowercase letters to the right of each bar indicate statistically similar samples via unpaired Student’s t-test (p > 0.05). (C) Representative images of indicated cells grown overnight at 25°C with brightfield and maximum intensity projections of Shs1-GFP in green and Elm1^KD^-mApple-GBP in magenta. Strains used are as follows from top to bottom: YEF9335 (*elm1^KD^-mApple-GBP*), YEF9362 (*SHS1-GFP elm1^KD^-mApple-GBP*), YEF10738 (*gin4Δ elm1^KD^-mApple-GBP*) and YEF10743 (*gin4Δ SHS1-GFP elm1^KD^-mApple-GBP*). Gray dashed line indicates the cell periphery. Scale bars = 5 μm. (D) Quantification of the ratio of round cells in strains used in Figures 7 A (Elm1^WT^) and 7 C (Elm1^KD^). Plotted is the average of 3 independent experiments of n ≥ 100 cells. Error bars are standard deviation. Lowercase letters to the right of each bar indicate statistically similar samples via unpaired Student’s t-test (p > 0.05).

### Gin4 anchors cortical septins to the plasma membrane in *elm1Δ* cells

One major difference in phenotypes between *elm1Δ* and *gin4Δ* cells is the prevalence and timing of the septins’ residency at the growing bud tip. Since the septins never had a sustained mislocalization to the growing bud cortex in any of the cells containing *gin4Δ* (**Fig. 3, A-D**) and Gin4 exhibits both potent septin- and plasma membrane -interacting capacity (Longtine et al., 1998a; Moravcevic et al., 2010; Mortensen et al., 2002), we wondered if the strong septin presence at the bud tip in *elm1Δ* cells could be due to the colocalized Gin4 there. To investigate this possibility, we first analyzed the septin localization behavior in the double deletion mutant *gin411 elm1Δ* cells. Surprisingly, while the cells still exhibited a strong elongated morphology phenotype typical of *elm1Δ* cells (**Fig. 8 A**), the septins behaved nearly identical with respect to kinetic timing to those in the *gin4Δ* single mutant (**Fig. 8, B and C**). A larger percentage of septin fluorescence left the bud neck in the double deletion mutant (more similar to the *elm1Δ* septin phenotype), but it quickly returned from the bud cortex after only 15-20 minutes of being displaced (**Fig. 8 C**).

**Figure 8.**
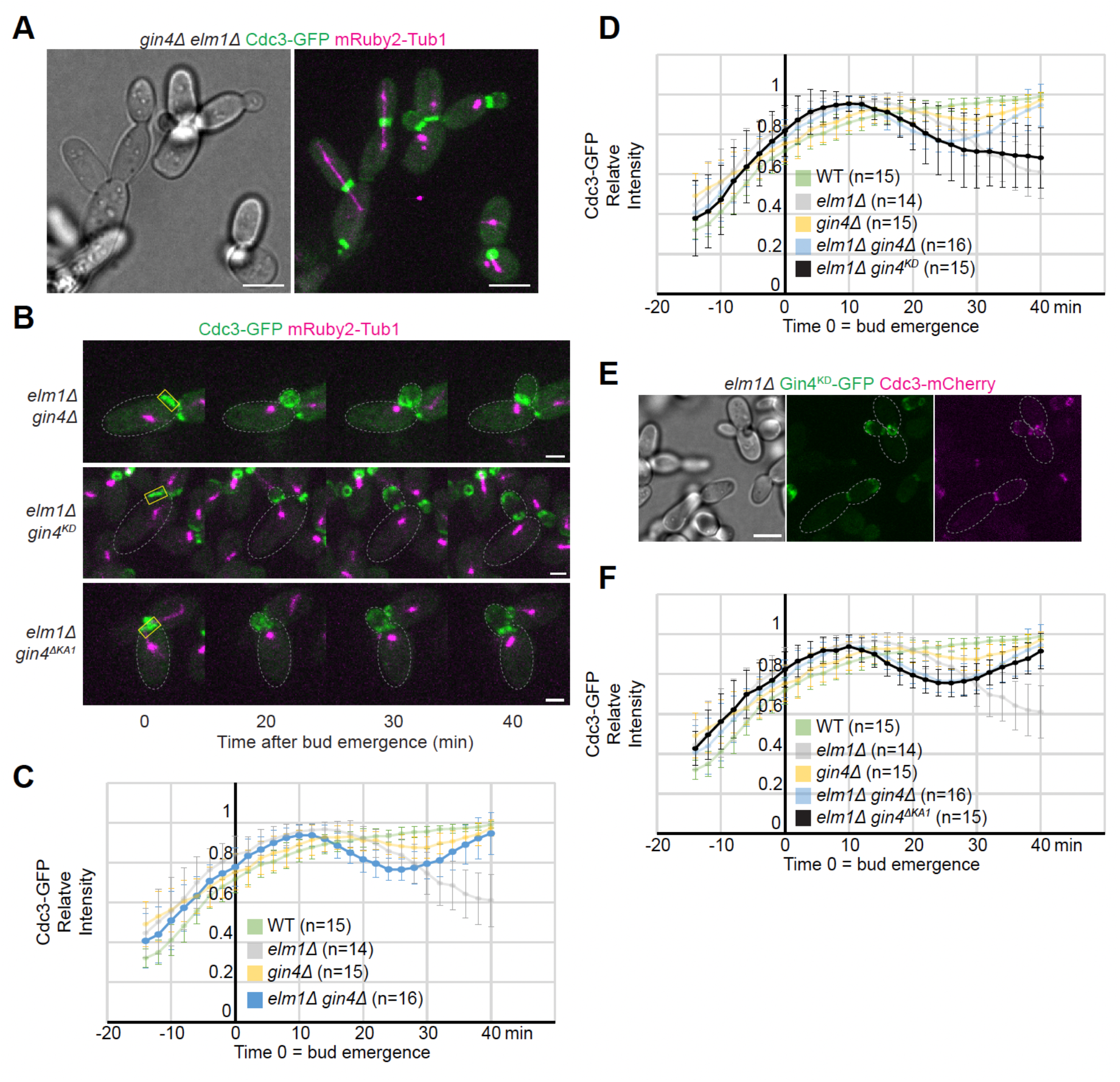
Gin4 retains septins at the bud cortex in *elm1Δ* cells. (A) Representative images of YEF10238 (*elm1Δ gin4Δ CDC3-GFP mRuby2-Tub1*) cells with brightfield (left) and maximum intensity projection of merged Cdc3-GFP in green and mRuby2-Tub1 in magenta (right). Scale bars = 5 μm. (B) Maximum-intensity projection images of representative *elm1Δ gin4Δ* (YEF10238), *elm1Δ gin4^KD^* (YEF11144), and *elm1Δ gin4^ΔKA1^* (YEF10990) cells from a time-lapse series taken with a 2-minute interval with Cdc3-GFP in green and mRuby2-Tub1 in magenta at indicated times. T = 0 is bud emergence. Gray dashed line is the cell periphery. Yellow boxed region is the area used for measurements in Figures 8 C, 8 D, and 8 F. Scale bars = 2 μm. (C) Quantification of cells in Figures 3 B and 8 B. Shown is background subtracted intensity of Cdc3-GFP in WT (green), *elm1Δ* (gray), *gin4Δ* (yellow), and *elm1Δ gin4Δ* (blue) relative to the maximum value measured from the sum projection of given number cells for each strain. Curves for WT, *elm1Δ*, and *gin4Δ* are same used in Figure 3 C and are at 30% opacity for ease of comparing kinetic signature of *elm1Δ gin4Δ* cells to WT and each single deletion mutant. The mean is plotted with error bars being the standard deviation. (D) Quantification of cells in Figures 3 B and 8 B. Shown is background subtracted intensity of Cdc3-GFP in WT (green), *elm1Δ* (gray), *gin4Δ* (yellow), *elm1Δ gin4Δ* (blue), and *elm1Δ gin4^KD^* (black) relative to the maximum value measured from the sum projection of given number cells for each strain. Curves for WT, *elm1Δ*, and *gin4Δ* are same used in Figure 3 C and that of *elm1Δ gin4Δ* is the same as used in Figure 8 C and are at 30% opacity for ease of comparing kinetic signature of *elm1Δ gin4^KD^* cells to previously analyzed cells. The mean is plotted with error bars being the standard deviation. (E) Representative image of YEF10989 (*elm1Δ gin4^KD^-GFP CDC3-mCherry*) cells with brightfield (left) and maximum intensity projection of Gin4^KD^-GFP in green (middle) and Cdc3-mCherry in magenta (right). Gray dashed line is cell periphery. Scale bar = 5 μm. (F) Quantification of cells in Figures 3 B and 8 B. Shown is background subtracted intensity of Cdc3-GFP in WT (green), *elm1Δ* (gray), *gin4Δ* (yellow), *elm1Δ gin4Δ* (blue), and *elm1Δ gin4^ΔKA1^* (black) relative to the maximum value measured from the sum projection of given number cells for each strain. Curves for WT, *elm1Δ*, and *gin4Δ* are same used in Figure 3 C and that of *elm1Δ gin4Δ* is the same as used in Figure 8 C and are at 30% opacity for ease of comparing kinetic signature of *elm1Δ gin4^ΔKA1^* cells to previously analyzed cells. The mean is plotted with error bars being the standard deviation.

Next, we addressed the question of whether this phenomenon was truly due to the presence of Gin4 or its lack of full activation by Elm1 (Asano et al., 2006) at the growing bud cortex in *elm1Δ* cells. In *elm1Δ* cells with the *gin4^KD^* allele at the endogenous locus, Cdc3-GFP maintained the same kinetic signature as that in the single *elm1Δ* strain (**Fig. 8, B and D**). Importantly, Gin4^KD^-GFP still exhibited bud cortex localization in *elm1Δ* cells (**Fig. 8 E**), presumably maintaining the bridge between the septins and the plasma membrane. In sharp contrast, Cdc3-GFP in *elm1Δ* Gin4^ΔKA1^ cells behaved nearly identical as in the *gin411 elm1Δ* double deletion mutant (**Fig. 8, B and F**). These data demonstrate that Gin4 is the membrane anchor that maintains septins at the growing bud cortex in *elmΔ* cells.

### Elm1 acts upstream of Gin4 to stabilize it at the division site before cytokinesis

While the above data indicate that Gin4 acts upstream of Elm1 to control its localization at the division site at the beginning of the cell cycle, previous work suggests that Elm1 most likely acts upstream of Gin4 during late mitosis (Asano et al., 2006; Sreenivasan and Kellogg, 1999); Mortensen, 2002 #2313}. To understand the underlying mechanism for this potential regulation, we first examined the localization kinetics of Gin4-GFP prior to cytokinesis in live cells. In *elm1Δ* cells, Gin4-GFP displaced from the bud neck region concurrently with the septins after bud emergence (**Fig. 2, A and B**). We have previously shown that the septins leave the bud cortex and return to the bud neck just prior to cytokinesis to participate in the HDR transition (Marquardt et al., 2020). In contrast, the amount of Gin4-GFP remaining at the bud neck continued to drop until only roughly 25% of the signal intensity remained when compared to that of WT cells and the signal never recovered prior to cytokinesis after it disappeared from the bud cortex (**Fig. 2 B and Fig. 9, A and B**). A more detailed investigation of the kinetics of Gin4-GFP just prior to cytokinesis in *elm1Δ* cells shows that the remaining Gin4-GFP signal began to dissociate form the bud neck 6-8 minutes earlier and with a much slower rate of loss than in WT cells (**Fig. 9, C and D**). These combined data, coupled with the observation that Gin4 started to dissociate after Elm1 was nearly gone from the bud neck (**Fig. 1 E and S1 D**), indicate that Elm1 most likely stabilizes Gin4 at the bud neck until the time for its removal in WT cells.

**Figure 9.**
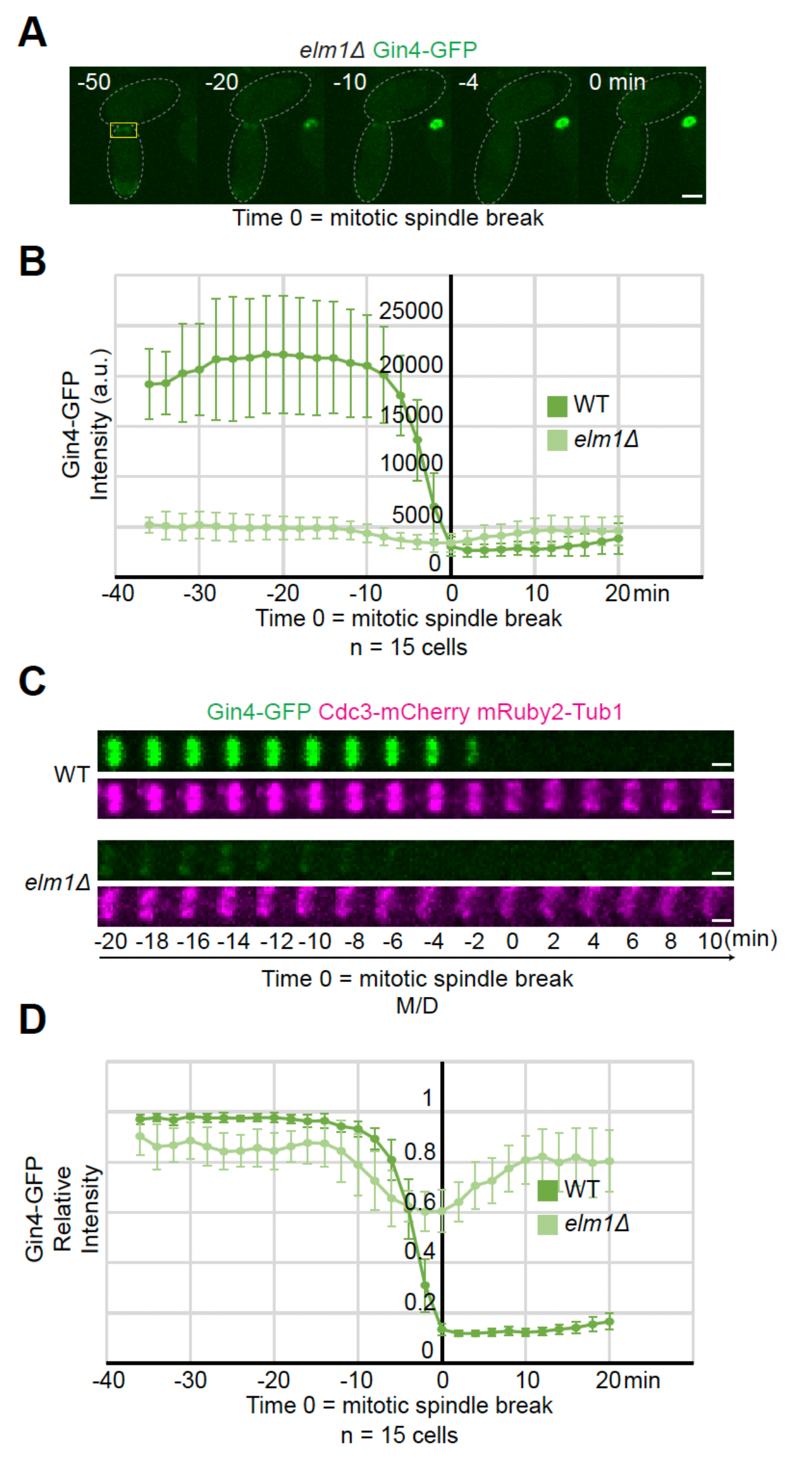
Elm1 stabilizes Gin4 at the bud neck prior to cytokinesis. (A) Maximum-intensity projection images of representative YEF10559 (*elm1Δ GIN4-GFP CDC3-mCherry mRuby2-Tub1*) cells from a time-lapse series taken with a 2-minute interval with Gin4-GFP in green and mRuby2-Tub1 in magenta at indicated times. T = 0 is mitotic spindle break. Gray dashed line is the cell periphery. Yellow boxed region is the area used for measurements in Figures 9 B and 9 D and to make montages for Figure 9 C. Scale bar = 2 μm. (B) Quantification of Gin4-GFP signal from cells in Figures 9 A and 9 C. Shown is integrated measured background subtracted intensity of Gin4-GFP in WT (dark green) and *elm1Δ* (light green) cells from the sum projection of given number cells for each strain. The mean is plotted with error bars being the standard deviation. A.U. = arbitrary units. (C) Montages of representative YEF10558 (*GIN4-GFP CDC3-mCherry mRuby2-Tub1*) and YEF10559 (*elm1Δ Gin4-GFP Cdc3-mCherry mRuby2-Tub1*) cells showing maximum-intensity projections from 20 minutes before to 10 minutes after mitotic spindle break from time-lapse series taken with a 2-minute interval. T = 0 is mitotic spindle break. Scale bars = 1 μm. (D) Quantification of Gin4-GFP signal from cells in Figures 9 A and 9 C. Shown is background subtracted intensity relative to the maximum value measured of Gin4-GFP in WT (dark green) and *elm1Δ* (light green) cells from the sum projection of given number cells for each strain. The mean is plotted with error bars being the standard deviation.

Elm1 was shown to directly phosphorylate Gin4 during mitosis (Asano et al., 2006). To determine if the phenotype we observed in *elm1Δ* cells with Gin4 showing decreased mitotic neck localization is due to phosphorylation, we examined the localization kinetics of Gin4 during late mitosis in cells with the *elm1^KD^* allele at the endogenous locus. These *elm1^KD^* cells exhibit a heterogenous phenotype with roughly 70% being elongated and the remaining 30% being round (Marquardt et al., 2020). We have previously shown that this phenotypic difference can be at least partially explained by the variable localization of Elm1^KD^ at the bud neck with round cells showing more localization and elongated cells showing little to no localization, and that artificially tethering Elm1^KD^ to the septin hourglass can largely rescue the elongated morphology (Marquardt et al., 2020). Gin4-mScarlet signal intensity was substantially lower prior to cytokinesis in both the elongated (51% when compared to WT cells) and round (66% when compared to WT cells) *elm1^KD^*cells, and it began to dissociate 30 minutes prior to cytokinesis in the elongated *elm1^KD^* cells and 25 minutes prior to cytokinesis in the round *elm1^KD^* cells, which was significantly altered from the 10 minutes prior to cytokinesis in WT cells (**Fig. 10, A-C**). The slight differences between the kinetic signatures of Gin4-mScarlet in WT cells when compared to our previous results using Gin4-GFP were most likely due to the photostability differences between the fluorescent tags used. We bypassed the phenotypic heterogeneity by again tethering various Elm1 mutants tagged with GBP at the genomic locus to the septin structure via Shs1-GFP. As expected, Elm1^WT^-GBP did not significantly alter the Gin4-mScarlet amount or kinetic signature prior to cytokinesis (**Fig. 10, D and E**). While Elm1^KD^-GBP did rescue the total signal intensity of Gin4-mScarlet at the bud neck, the Gin4-mScarlet signal began to dissociate almost 10 minutes earlier than in either the WT or Elm1^WT^-GBP cells (**Fig. 10, D and E**). This indicates that Elm1 phosphorylation of Gin4 during late mitosis most likely maintains Gin4 stability at the bud neck.

**Figure 10.**
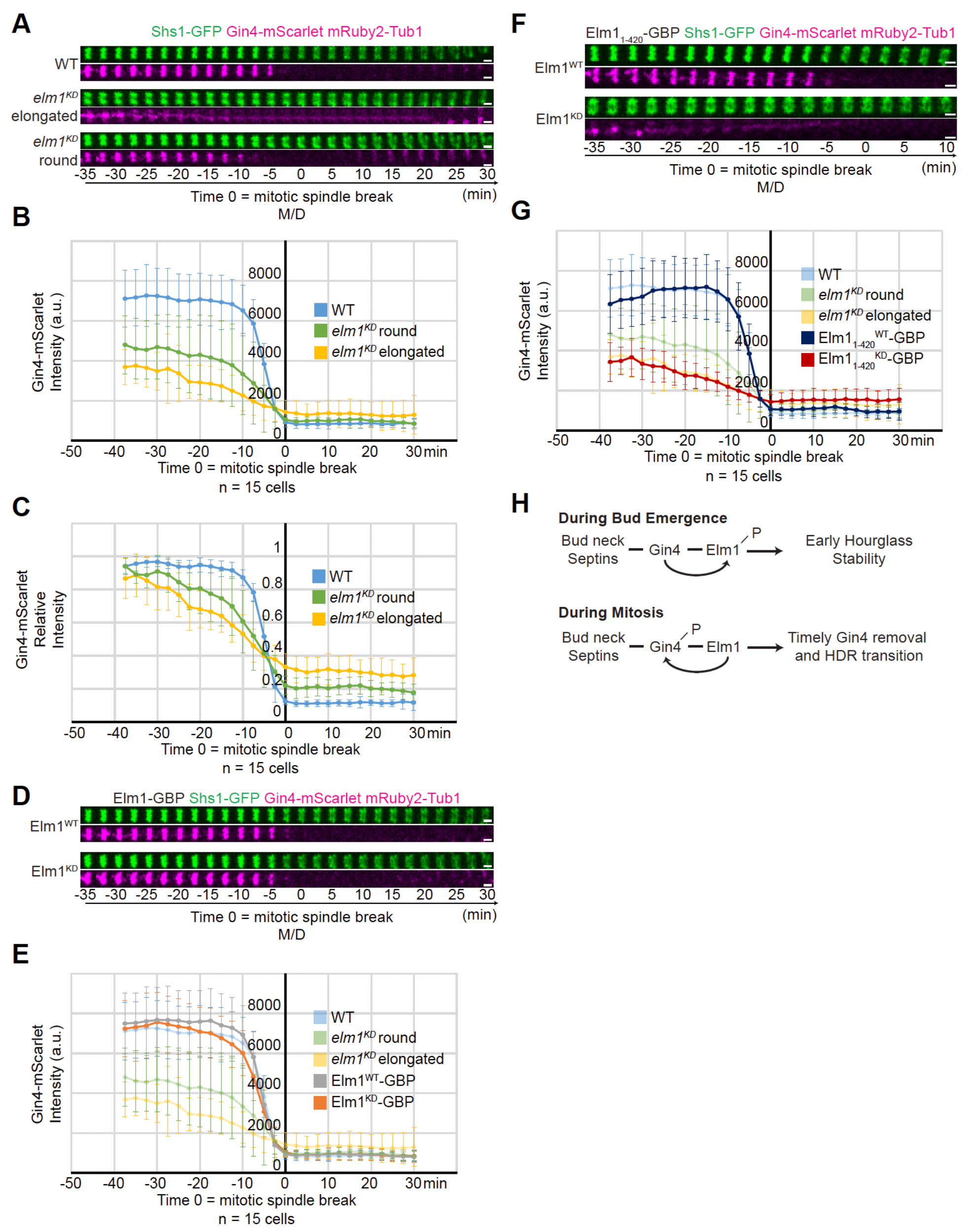
Elm1 binding and kinase activity are required for the retention of Gin4 at the bud neck prior to cytokinesis. (A) Montages of bud neck region of representative WT (top, Strain YEF11493), elongated *elm1^KD^*(middle, Strain YEF11496), and round *elm1^KD^* (bottom, Strain YEF11496) cells showing maximum-intensity projections of Shs1-GFP in green and Gin4-mScarlet mRuby2-Tub1 in magenta from 35 minutes before to 30 minutes after mitotic spindle break from time-lapse series taken with a 2.5-minute interval. T = 0 is mitotic spindle break. Scale bars = 1 μm. (B) Quantification of Gin4-mScarlet signal from cells in Figure 10 A. Shown is integrated measured background subtracted intensity of Gin4-mScarlet in WT (blue), round *elm1^KD^* (green), and elongated *elm1^KD^* (yellow) cells from the sum projection of given number cells for each strain. The mean is plotted with error bars being the standard deviation. A.U. = arbitrary units. (C) Quantification of Gin4-mScarlet signal from cells in Figure 10 A. Shown is background subtracted intensity relative to the maximum value measured of Gin4-mScarlet in WT (blue), round *elm1^KD^* (green), and elongated *elm1^KD^* (yellow) cells from the sum projection of given number cells for each strain. The mean is plotted with error bars being the standard deviation. (D) Montages of bud neck region of representative Elm1^WT^-GBP (top, Strain YEF11495) and Elm1^KD^-GBP (bottom, Strain YEF11494) cells showing maximum-intensity projections of Shs1-GFP in green and Gin4-mScarlet mRuby2-Tub1 in magenta from 35 minutes before to 30 minutes after mitotic spindle break from time-lapse series taken with a 2.5-minute interval. T = 0 is mitotic spindle break. Scale bars = 1 μm. (E) Quantification of Gin4-mScarlet signal from cells in Figures 10 A and 10 D. Shown is integrated measured background subtracted intensity of Gin4-mScarlet in WT (blue), round *elm1^KD^* (green), elongated *elm1^KD^*(yellow), *elm1^WT^-GBP* (gray), and *elm1^KD^-GBP* (orange) cells from the sum projection of given number cells for each strain. Curves for WT, round *elm1^KD^*, and elongated *elm1^KD^*are same used in Figure 10 C and are at 30% opacity for ease of comparing kinetic signature of *elm1^WT/KD^-GBP* cells to previously analyzed cells. The mean is plotted with error bars being the standard deviation. A.U. = arbitrary units. (F) Montages of bud neck region of representative *elm1_1-420_^WT^-GBP* (top, Strain YEF11607) and *elm1_1-420_^KD^-GBP* (bottom, Strain YEF11606) cells showing maximum-intensity projections of Shs1-GFP in green and Gin4-mScarlet mRuby2-Tub1 in magenta from 35 minutes before to 30 minutes after mitotic spindle break from time-lapse series taken with a 2.5-minute interval. T = 0 is mitotic spindle break. Scale bars = 1 μm. (G) Quantification of Gin4-mScarlet signal from cells in Figures 10 A and 10 F. Shown is integrated measured background subtracted intensity of Gin4-mScarlet in WT (blue), round *elm1^KD^* (green), elongated *elm1^KD^* (yellow), *elm1_1-420_^WT^-GBP* (navy blue), and *elm1_1-420_^KD^-GBP* (red) cells from the sum projection of given number cells for each strain. Curves for WT, round *elm1^KD^*, and elongated *elm1^KD^*are same used in Figure 10 C and are at 30% opacity for ease of comparing kinetic signature of *elm1_1-420_^WT/KD^-GBP* cells to previously analyzed cells. The mean is plotted with error bars being the standard deviation. A.U. = arbitrary units. (H) Model depiction of Gin4 and Elm1 synergistic regulation at bud emergence (top) and prior to cytokinesis (bottom). Solid lines indicate direct interacting partners, arrows indicate direction of regulation by binding and phosphorylation (denoted by a P).

Since Elm1 and Gin4 interact with each other and this interaction is critical for Elm1 localization and function during the septin hourglass stage **(Fig. 4 and 5),** we next investigated if the same is true for Gin4 localization and function prior to cytokinesis. The Elm1 kinase domain (Elm1_1-420_) did not interact with Gin4 in vitro (**Fig. 4 B**). By tagging this fragment with GBP at the endogenous locus to bypass the localization loss (Marquardt et al., 2020; Moore et al., 2010) and tethering it to the septins via Shs1-GFP, we could investigate the necessity of the Elm1-Gin4 interaction prior to cytokinesis. Tethering the WT kinase domain (Elm1_1-420_^WT^-GBP) to Shs1-GFP resulted in a complete recovery of both the neck localization of Gin4-mScarlet and its steep removal 10 minutes prior to cytokinesis (**Fig. 10, F and G**). However, the kinase-dead variant (Elm1_1-420_^KD^-GBP) failed to rescue either aspect of Gin4 function and the Gin4-mScarlet kinetic profile most resembled that of untethered *elm1^KD^* cells (**Fig. 10, F and G**). This allows us to conclude that Elm1 binding and phosphorylation of Gin4 are both critical for maintaining Gin4 at the bud neck during the late hourglass stage prior to cytokinesis, similar to the regulation of Gin4 on Elm1 during the early hourglass stage.

## DISCUSSION

### Mutual control and functional impact between the septin hourglass-associated kinases Elm1 and Gin4 during the cell cycle

Mechanistic analysis of septin high-order assembly and architectural remodeling has been an active area of research since septin filaments were first visualized in the 1970s (Byers and Goetsch, 1976). While many details in the budding yeast *S. cerevisiae* such as septin octamer composition and filament organization have been illustrated in recent years (Bertin et al., 2008; Bertin et al., 2012; Bertin and Nogales, 2016; Chen et al., 2020; DeMay et al., 2011; Garcia et al., 2011; Ong et al., 2014; Rodal et al., 2005; Weems and McMurray, 2017), the regulations involved at each step by SAPs remain largely unanswered. The protein kinase Elm1 controls septin hourglass assembly and stability at the division site at least in part by regulating septin filament pairing (Marquardt et al., 2020). Elm1 also regulates many other processes in *S. cerevisiae* including cellular morphogenesis, the spindle position checkpoint, and metabolism (Caydasi et al., 2010; Kang et al., 2016; Moore et al., 2010; Sutherland et al., 2003). However, the regulation of Elm1 itself has not been elucidated. Here we have shown that a second septin hourglass-associated kinase, the Nim1-related kinase Gin4, is responsible for the proper targeting of Elm1 to the septin hourglass structure and its maintenance there through a combination of direct binding and phosphorylation (**Fig. 10 H**, top). Elm1 is not known to have any direct binding to the septin structure. The only other SAP with which it is known to have a strong direct interaction is Bni5 (Marquardt et al., 2020; Patasi et al., 2015); however, deletion of *BNI5* does not affect the bud neck localization of Elm1 in otherwise WT cells (Marquardt et al., 2020). Thus, our discovery of the Gin4-dependent localization represents a significant finding in Elm1 regulation.

We also found that Gin4 phosphorylates Elm1 on at least seven residues mostly located in the C-terminal bud neck-localization domain. These phosphorylation events are critical for Elm1 localization to the bud neck during the hourglass stage (**Fig. 5, C and D**). However, Gin4 may lose affinity for Elm1 after the phosphorylation has happened, because the Elm1^7D^ mutant protein fails to interact with Gin4 in vitro to the same degree as the other mutant proteins (**Fig. 5, E and F**). Kinases tend to have transient interactions with their substrates to encourage additional rounds of phosphorylation (Niinae et al., 2021), therefore this loss in binding affinity is not surprising. It appears that Elm1^7D^ requires the presence of Bni5 at the bud neck for its residual localization at the bud neck (**Fig. S4**), which implies that Gin4 may recruit Elm1 to the bud neck, phosphorylate it, and then pass it along to interact with Bni5. Elm1 and Bni5 interact in vitro independent of Gin4 (Marquardt et al., 2020), thus, the Elm1 phosphorylation by Gin4 is not required or is bypassed for its interaction with Bni5 in vitro. Nonetheless, this analysis reveals an elaborate control of Elm1 localization and interaction at the bud neck by Gin4 at the early stage of the cell cycle.

The septin and cell shape phenotypes of *elm1Δ* cells are much more pronounced than those of *gin4Δ* cells (**Fig. 3**) (Gladfelter et al., 2004). It is therefore surprising that Gin4 acts upstream of Elm1 at the septin hourglass stage. However, two important pieces of data generated in this study can explain this phenomenon. Firstly, Gin4 interaction with and phosphorylation of Elm1 does not seem to alter the kinase activity of Elm1. Bni5 is essential for normal septin and cell morphology in the presence of an *elm1^KD^* allele (Marquardt et al., 2020), yet deletion of *BNI5* does not exacerbate any observed phenotypes associated with the phospho-mutant alleles of *ELM1* at the endogenous locus (**Fig. S4**). Thus, the cytoplasmic phospho-mutants of Elm1 must retain kinase activity. When the kinase domain alone is expressed and absent from the bud neck, only a proportion of cells exhibit a septin and cell shape phenotype that is exacerbated if the dead kinase domain is expressed (Marquardt et al., 2020). Thus, the lack of interaction with Gin4 does not appear to compromise the kinase activity of Elm1. Secondly, Gin4 anchors the septins to the growing bud tip in *elm1Δ* cells (**Fig. 8**). If the septins were able to relocate from the growing tip to the bud neck in *gin4Δ* cells in a timely fashion, it could simply be a “delay” in hourglass assembly that the cells can tolerate and appear relatively normal later in mitosis. Collectively, our results indicate that the elaborate control of Elm1 localization and function by Gin4 at the beginning of the cell cycle plays a major role in controlling septin hourglass assembly and stability.

Elm1 is known to regulate Gin4 during mitosis through direct phosphorylation (Asano et al., 2006). Here we have extended this regulation to also include direct binding of Elm1 to Gin4 which is of critical importance in maintaining an adequate bud-neck level of Gin4 prior to cytokinesis (**Fig. 10 H**, bottom). We also discovered that some of the sites, originally thought to be Elm1 phosphorylation sites in Gin4 (Asano et al., 2006), were phosphorylated in our in vitro kinase reaction even though the only Elm1 species is a dead kinase (**Fig. S2 C**, asterisks). This may indicate that either our Elm1^KD^ is not completely inactive or that the sites are Gin4 auto-phosphorylation sites when Gin4 is bound to Elm1. Indeed, Gin4^KD^ has been shown to exhibit auto-phosphorylation when visualized by autoradiography (Asano et al., 2006), leading us to lean toward the latter possibility. Since deletion of either *ELM1* (Marquardt et al., 2020) or *GIN4* (**Fig. 3, F - H**) affected the HDR transition presumably through their roles in septin hourglass assembly and maturation, the stabilization of Gin4 at the bud neck by Elm1 during mitosis is likely involved in controlling the timely HDR transition. The finding that septins in both *elm1Δ* and *gin4Δ* cells showed a lower intensity at the bud neck before the HDR transition, and a lower loss of intensity and altered kinetics during the transition suggests that the septins may reorganize with the removal of Gin4 protein from the bud neck just prior to cytokinesis. Gin4 has potent septin- and plasma membrane binding capacity (Longtine et al., 1998a; Moravcevic et al., 2010; Mortensen et al., 2002) and departs the bud neck with similar kinetic timing to that of the loss of signal of Cdc3 (**Fig. 1 E**). Gin4 levels at the bud neck just prior to cytokinesis are substantially lower in *elm1Δ* cells and depart earlier than that of WT (**Fig. 9**), which is similar to the kinetic timing of septins in *elm1Δ* cells (**Fig. 3, F and G**). All of these data lead us to conclude that the Elm1-controlled removal of Gin4 in mitosis promotes timely HDR transition at least in part by promoting timely removal of the hourglass septins during the reorganization.

The finding that both kinases utilize similar strategies for regulation (direct binding for appropriate targeting and direct phosphorylation for protein stability at the bud neck) illustrates that the cell is able to accomplish two functional tasks with the same protein machinery, which defines an elegant mechanism for the mutual control of Elm1 and Gin4 in septin hourglass assembly and remodeling during the cell cycle.

### Shared and unique roles of Elm1 and Gin4 in septin hourglass assembly and function and beyond

Besides their shared role in controlling septin hourglass assembly, both Elm1 and Gin4 are also required for the localization of the formin Bnr1 to the division site before cytokinesis (Buttery et al., 2012). While Gin4 interacts directly with Bnr1, the mechanism for the role of Elm1 in this process remains unknown (Buttery et al., 2012). It will be important to determine whether Elm1 and Gin4 mutually regulate in the same pathway to control Bnr1 localization and whether their roles in this process depend on their roles in the control of septin hourglass assembly.

The observations that Elm1-tethering to the bud neck independently of Gin4 cannot fully rescue all the phenotypes of *gin4Δ* cells (**Fig. 7**) and that the double deletion of *ELM1* and *GIN4* behave as a combination of the two deletions (**Fig. 8**) suggest that while these two kinases share functions by operating in a single regulatory pathway, each kinase must also possess some unique functionality. Gin4 exerts additional septin regulation possibly through direct phosphorylation of the septin Shs1 to potentially regulate the HDR (Asano et al., 2006; Mortensen et al., 2002), whereas Elm1 exhibits direct regulation of the morphogenesis checkpoint by phosphorylating the kinase Hsl1 to initiate the sequestering of the CDK inhibitor Swe1 to the bud neck to allow mitotic progression (Kang et al., 2016; Szkotnicki et al., 2008). It will be important to further investigate these regulatory networks to determine whether any synergistic relationships exist between Elm1 and Gin4.

Beyond the *S. cerevisiae* model system, little is known on the direct synergistic roles of Elm1 and Gin4 functional orthologs. However, it is noteworthy that deletion of the *ELM1* or *GIN4* ortholog in the filamentous fungus *Ashbya gossypii* specifically abolishes the formation of the inter-region (IR) septin rings, not the septin structures near the hyphal tips or at the branch sites (DeMay et al., 2009). Importantly, septin filaments in the IR rings are organized in parallel to the long hyphal axis, similar to the paired filaments in the septin hourglass in budding yeast (DeMay et al., 2011; Ong et al., 2014). These observations raise the possibility that the Elm1 and Gin4 orthologs in *A. gossypii* may act in the same pathway as seen in budding yeast to control the IR septin ring assembly.

During *Caenorhabditis elegans* embryogenesis, both the Elm1 (PAR-4)- and Gin4 (PAR-1)-like proteins are involved in initial polarity establishment (Guo and Kemphues, 1995; Watts et al., 2000). Unlike in *S. cerevisiae*, *par-1* mutants do not affect the localization of the PAR-4 protein distribution (Watts et al., 2000), but it is believed that they may exist in the same (or at least a partially overlapping) genetic pathway due to similar phenotypes upon their inactivation (Watts et al., 2000). Similar data was obtained in *Drosophila melanogaster* since both PAR-1 and LKB1 regulate anterior-posterior polarity in oocytes (Martin and St Johnston, 2003; Shulman et al., 2000; Tomancak et al., 2000). Importantly, PAR-1 was shown to phosphorylate LKB1 in vitro (Martin and St Johnston, 2003). As the C-terminal non-kinase domain of *S. cerevisiae* Elm1 (where most of the direct phosphorylation was discovered in our study) is not well conserved outside of fungal species, the PAR-1-catalyzed phosphorylation sites in LKB1 (not yet identified) are unlikely to be the same as those Gin4-catalyzed sites in Elm1. However, this does not rule out the possibility that the interplay between PAR-4/LKB1 and PAR-1 in polarity axis formation during embryogenesis in *C. elegans* and Drosophila is mechanistically or conceptually similar to the interplay between Elm1 and Gin4 in controlling septin hourglass assembly (this study) and/or polarized cell growth in budding (Carroll et al., 1998; Sreenivasan and Kellogg, 1999; Tjandra et al., 1998).

## MATERIALS AND METHODS

### Stains and manipulations

The budding yeast *Saccharomyces cerevisiae* is our experimental model. All yeast strains used in this study are in the YEF473A background (Bi and Pringle, 1996) and are listed in **Table S1**. Standard culture media and genetic techniques were used (Guthrie and Fink, 1991). Yeast strains were grown routinely at 25°C in synthetic complete (SC) minimal medium lacking specific amino acid(s) and/or uracil or in rich medium YM-1 (Lillie and Pringle, 1980) or yeast extract/peptone/dextrose (YPD). New strains were constructed either by integrating a plasmid carrying a modified gene at a genomic locus or by transferring a deletion or tagged allele of a gene from a plasmid or from one strain to another via PCR amplification and yeast transformation (Lee et al., 2013; Longtine et al., 1998b) (see footnotes in **Table S1**).

For spot growth assay in Fig. 6 A and Fig. S4 A, ten-fold serial dilutions of cell cultures of the indicated strains were spotted on SC plates and incubated at 25°C or 37°C for 3 days prior to image capture and analysis.

### Oligonucleotides and plasmids

All primers were purchased from Integrated DNA Technologies and are written 5’-3’ for all sequences below. Primers used for deletions of genes are as follows: *ELM1*, Elm1-F-check (GAGGAACTTACTTGATCCTTCTTGAAG) and Elm1-R-check (GATTTCGCGACACAGTGG); *GIN4*, Gin4-F1 (AGAAAGATATTCGCAGCACAATACAATAATAACATTCAAACGGATCCCCGGGTTAATTAA), Gin4-R1 (CAAAACGAAGGAGACAAAACATGATTGCATTACATTAGCAGAATTCGAGCTCGTTTAAAC), Gin4-F-check (CACTTCACTGGAAGAACTGGG), and Gin4-R-check (GCTCTTACTTTAATCCCAAAGAGG); *SHS1*, Shs1-F-check (ACCACCTTTTTCCATACGA) and Shs1-R-check (GTTACGGGAAATCATGATAG); *BNI5*, F-BNI5-370up-ATG (AAGAGATTGACGGTGGATGGAGAC) and R-BNI5-510down-TGA (TGTGCAGTTTGTTTCGAAAACCATC). Primers used for C-terminal tagging at the genomic locus are as follows: *ELM1,* Elm1-Ftag-check (CCTAAAGAGAACGGGAACAGAAC) and Elm1-R-check; *GIN4*, Gin4-F5 (AGTTGAGAACGTCCTGAATAAGGAAGGCGTTCTACAAAAATGGTGACGGTGCTGGTTTA), Gin4-R3 (CAAAACGAAGGAGACAAAACATGATTGCATTACATTAGCATCGATGAATTCGAGCTCG), Gin4-Ftag-check (GGAATTGTATGCCAAGATTTCTG), and Gin4-R-check; *SHS1*, Shs1-Ftag-check (GAAACCGTTCCATATGTCTTG) and Shs1-R-check. Primers used for N-terminal tagging of the endogenous *ELM1* locus were F-Elm1-prom-GFP (TTTTTTGAACGCCAGGTTAACAATAATTACTTAGCATGAAATGTCTAAAGGTGAAGAATT) and R-Elm1-term-Elm1 (640) (CAGCTAACCCAATCCGACAGATATCATCCTGTAGTTTCATCTATATTTGACCATTATCTG). Primers used to delete the KA1 domain from the endogenous *GIN4* locus were Gin4-F-check and Gin4-dKA1-R (AACGAAGGAGACAAAACATGATTGCATTACATTAGCACTATGCCTGCGAGCCAGCATTTT). Primers used to mutagenize lysine 48 to alanine in the *GIN4* locus were Gin4-K48A-F (CAGGACAAGAGGCGGCAGTTGCGGTAATATCAAAAGCAGTATT) and Gin4-K48A-R (AATACTGCTTTTGATATTACCGCAACTGCCGCCTCTTGTCCTG). Primers used to mutagenize lysine 117 to arginine in the *ELM1* locus were Elm1-K117R SDM F (TAGGCAAGGTTGTTGCTGTCAGGATTATACCAAAAAAACCTTG) and Elm1-K117R SDM R (CAAGGTTTTTTTGGTATAATCCTGACAGCAACAACCTTGCCTA). All site-directed mutagenesis listed above was confirmed correct via sequencing at the DNA Sequencing Facility, University of Pennsylvania.

Plasmids YIp128-CDC3-GFP (Caviston et al., 2003) and YIp128-CDC3-mCherry (Gao et al., 2007) (integrative, LEU2) carry an N-terminally GFP- or mCherry-tagged CDC3 under the control of its own promoter, respectively. Plasmids pHIS3p::mRuby2-Tub1+3’UTR::HPH (Markus et al., 2015) and pFA6a-link-ymScarlet-I-CaURA, pGEX-4T1-Elm1FL, pGEX-4T1-Elm1_1-420_, and pGEX-4T1-Elm1_421-640_ (Marquardt et al., 2020) were described previously. Plasmid pFA6a-URA3-KanMX6 was a generous gift from John Pringle and is described previously (Onishi et al., 2013). Plasmid pGEX-4T1 was purchased from GE Healthcare (Cat #: 28-9545-49). Plasmid pET His6 Sumo TEV LIC (Addgene # 29659) was a generous gift from Scott Gradia.

The following plasmids were generated for this study: pGEX-4T1-Elm1^KD^ ^(K117R)^, pET-His6-Sumo-Gin4, pET-His6-Sumo-Gin4^KD^ ^(K48A)^, pUG36-Elm1, pUG36-Elm1^S519A^, pUG36-Elm1^S519D^, pUG36-Elm1^7A^, pUG36-Elm1^7D^, pGEX-4T1-Elm1^S519A^, pGEX-4T1-Elm1^S519D^, pGEX-4T1-Elm1^7A^, and pGEX-4T1-Elm1^7D^. To generate pGEX-4T1-Elm1^KD(K117R)^, mutagenesis primers Elm1-K117R SDM F and Elm1-K117R SDM R were used to amplify pGEX-4T1-Elm1, and the parental plasmid was digested with DpnI (New England Biolabs, Ipswitch, MA, USA) prior to transformation into *E. coli*. To generate pET-His6-Sumo-Gin4, *GIN4* was PCR amplified using genomic DNA from YEF473A as the template DNA using primers Gin4 FL-SspI (TGTACTTCCAATCCAATATTATGGCAATCAATGGTAACAG) and Gin4 FL-BamHI (CGGCGCTCGAATTCGGATCCCTATTTTTGTAGAACGCCTTC), and subsequently digested with SspI and BamHI (New England Biolabs) before being ligated into pET-His6-Sumo digested with the same enzymes. To generate pET-His6-Sumo-Gin4^KD(K48A)^, mutagenesis primers Gin4-K48A-F and Gin4-K48A-R were used to amplify pET-His6-Sumo-Gin4, and the parental plasmid was digested with DpnI prior to transformation into *E. coli*. To generate pUG36-Elm1, *ELM1* was PCR amplified using genomic DNA from YEF473A as the template DNA using primers F-pUG36-link-ELM1-start (GTACAAATCTAGAACTAGTGGATCCCCCGGGCTGCAGATGTCACCTCGACAGCTTATACC) and R-pUG36-link-ELM1-Stop (ATGACTCGAGGTCGACGGTATCGATAAGCTTGATATCCTATATTTGACCATTATCTGCAA), and was transformed into YEF473A with EcoRI (New England Biolabs)-digested pUG36 (a kind gift from Johannes H. Hegemann) and repaired by gap-repair cloning. To generate pUG36-Elm1^S519A^ and pUG36-Elm1^S519D^, mutagenesis primers Elm1 S519A-F (CTTTTTGTAGGTCAAATGAAGCCTTACCTAATTTGACTGTGAA) and Elm1 S519A-R (TTCACAGTCAAATTAGGTAAGGCTTCATTTGACCTACAAAAAG) or Elm1 S519D-F (CTTTTTGTAGGTCAAATGAAGACTTACCTAATTTGACTGTGAA) and Elm1 S519D-R (TTCACAGTCAAATTAGGTAAGTCTTCATTTGACCTACAAAAAG), respectively, and the parental plasmid digested with DpnI before being transformed into *E. coli*. To generate pUG36-Elm1^7A^, mutagenesis of each phosphorylation site was achieved with subsequent rounds of mutagenesis PCR followed by transformation into *E. coli*. First, S604A and S605A was introduced to pUG36-Elm1^S519A^ using primers Elm1-S604A S605A-F (GGGCTAGAAAGCTTGCTCATGCAGCTAATATTCTCAACTTTAAAGC) and Elm1-S604A S605A-R (GCTTTAAAGTTGAGAATATTAGCTGCATGAGCAAGCTTTCTAGCCC); this plasmid was then used as a template to mutagenize S324A using primers Elm1-S324A-F (AGCTATGTCATTTGGGCAATGCCAAAAGAGATTTTGTGACGGA) and Elm1-S324A-R (TCCGTCACAAAATCTCTTTTGGCATTGCCCAAATGACATAGCT); this plasmid was then used as a template to mutagenize S422A and S424A using primers Elm1-S422A S424A-F (GGAATCACAGTCAAATTTCAGCGTCCGCTGTGAACCCCGTAAGAAACGA) and Elm1-S422A S424A-R (TCGTTTCTTACGGGGTTCACAGCGGACGCTGAAATTTGACTGTGATTCC); lastly, this plasmid was used as a template to mutagenize T463A using primers Elm1-T463A-F (ACAAGGTATTGGTATCTGCAGCTAGTAAAGTAACACCTTCGAT) and Elm1-T463A-R (ATCGAAGGTGTTACTTTACTAGCTGCAGATACCAATACCTTGT). To generate pUG36-Elm1^7D^, the same strategy was employed except the following primers were used to make S/T-D mutagenesis: Elm1-S604D S605D-F (GGGCTAGAAAGCTTGCTCATGACGATAATATTCTCAACTTTAAAGC) and Elm1-S604D S605D-R (GCTTTAAAGTTGAGAATATTATCGTCATGAGCAAGCTTTCTAGCCC), Elm1-S324D-F (AGCTATGTCATTTGGGCAATGACAAAAGAGATTTTGTGACGGA) and Elm1-S324D-R (TCCGTCACAAAATCTCTTTTGTCATTGCCCAAATGACATAGCT), Elm1-S422D S424D-F (GGAATCACAGTCAAATTTCAGACTCCGATGTGAACCCCGTAAGAAACGA) and Elm1-S422D S424D-R (TCGTTTCTTACGGGGTTCACATCGGAGTCTGAAATTTGACTGTGATTCC), and Elm1-T463D-F (ACAAGGTATTGGTATCTGCAGATAGTAAAGTAACACCTTCGAT) and Elm1-T463D-R (ATCGAAGGTGTTACTTTACTATCTGCAGATACCAATACCTTGT). To generate pGEX-4T1-Elm1^S519A^, pGEX-4T1-Elm1^S519D^, pGEX-4T1-Elm1^7A^, and pGEX-4T1-Elm1^7D^, pUG36-Elm1^S519A^, pUG36-Elm1^S519D^, pUG36-Elm1^7A^, pUG36-Elm1^7D^ were used as template DNA, respectively using the primers GST-Elm1-F (GGTTCCGCGTGGATCCATGTCACCTCGACAGCTTATACCG) and GST-Elm1-R (GATGCGGCCGCTCGAGCTATATTTGACCATTATCTGCAAAG) to amplify each Elm1 phospho-mutant and cloned into pGEX-4T1 using BamHI and XhoI (New England Biolabs). All plasmid constructs and mutagenesis were confirmed correct via sequencing at the DNA Sequencing Facility, University of Pennsylvania.

### Live cell imaging and quantitative analysis

For time-lapse microscopy, cells were grown at 25°C to exponential phase in liquid SC media. Cells were briefly sonicated (model Q55, Qsonica, Newtown, CT, USA) at 15% power for 5 seconds to declump, concentrated by centrifugation, spotted onto a poly-lysine-coated glass-bottom dish, and then embedded with SC containing agarose (Okada et al., 2017) or spotted onto a concanavalin A (Sigma)-coated glass-bottom dish, and then SC medium was added (Okada et al., 2021). Images were acquired at room temperature using a Nikon microscope (model Eclipse Ti-U, Tokyo, Japan) equipped with a Nikon 100x/1.49NA oil objective (model CFI Apo TIRF 100x), and a Yokogawa spinning-disk confocal scanner unit (model CSU-X1, Tokyo, Japan). A Photometrics Evolve Delta 512X512 EMCCD Digital Monochrome Camera (Tucson, AZ, USA) was used for image capture. Solid-state lasers for excitation (488 nm for GFP and 561 nm for RFP) were housed in a launch constructed by Spectral Applied Research (model ILE-400, Richmond Hill, Ontario, Canada). The imaging system was controlled by MetaMorph version 7.8.10.0 (Molecular Devices, Downingtown, PA, USA). Images were taken with z-stacks of 13 × 0.7 μm for all the experiments.

For quantification of the fluorescence intensity of targeted protein signal (Elm1-GFP, Gin4-GFP, Cdc3-GFP, Cdc3-mCherry, and Gin4-mScarlet), a sum projection was created, and quantification was performed with ImageJ (National Institutes of Health). A polygon was drawn around the region of interest and the integrated density was measured along the entire time-lapse series. The fluorescence intensity of the background was subtracted from this measurement (outside the cell). For Figures 2 E, 4 D, 4 E, 9 B, 10 B, 10 E, 10 G, S3 B (top), and S3 D (top) this value was used for data analysis. For Figures S3 B (bottom) and S3 D (bottom) the same method was used, but only time point 20 minutes after bud emergence was used for analysis. For Figure 3 H, the measured intensity data was normalized for region of interest area to directly compare the density of the signal intensity per unit area since measured regions had a vast size discrepancy between and among the different mutant strains. For Figures 1 C, 1 E, 2 B, 2 D, 3 C, 3G, 8 C, 8 D, 8 F, 9 D, 10 C, S1 B and S1 D, the measured intensity data were normalized to the peak intensity (100%) of the fusion protein signal at the bud neck in the cell. Data were analyzed with Microsoft Excel and expressed as the mean value ± standard deviation (SD) except for Figures S3 B (bottom) and S3 D (bottom) in which each cell’s measured intensity is plotted with the black line indicating the sample’s average value. For Figure 6 D, septin phenotypes were analyzed from time-lapse series and categorized as either septin signal in the growing bud cortex or bud-neck septins asymmetrically localized to the daughter side of the bud neck and plotted as a ratio to the total number of cells with each indicated phenotype.

For snapshot images, cells were grown at 25°C or 37°C to exponential phase in liquid SC media. Cells were briefly sonicated at 15% power for 5 seconds to declump, concentrated by centrifugation, and spotted onto a slide. Images were taken as described above for all the experiments. To quantify cell numbers with elongated phenotypes (**Fig. 7, B and D**), cells were counted as round if the bud length: width ≤1.2, while they were counted as elongated if >1.2.

### Protein purification and in vitro binding assays

Rosetta DE3 cells transformed with either pGEX-4T1 (GST alone), pGEX-4T1-Elm1^WT^ (GST-Elm1^WT^), pGEX-4T1-Elm1^KD^ (GST-Elm1^KD^), pGEX-4T1-Elm1_1-420_ (GST-Elm1_1-420_), pGEX-4T1-Elm1_421-640_ (GST-Elm1_421-640_), pGEX-4T1-Elm1^S519A^ (GST-Elm1^S519A^), pGEX-4T1-Elm1^S519D^ (GST-Elm1^S519D^), pGEX-4T1-Elm1^7A^ (GST-Elm1 Elm1^7A^), pGEX-4T1-Elm1^7D^ (GST-Elm1 Elm1^7D^), pET-His6 Sumo-TEV-LIC (6xHis-SUMO alone), pET-His6-Sumo-Gin4 (6xHis-SUMO-Gin4), or pET-His6-Sumo-Gin4^KD^ (6xHis-SUMO-Gin4^KD^) were grown to an OD600 0.6-1.0 before being induced for 3 hours with 0.3mM IPTG (Lab Scientific, Highlands, NJ, USA) at 25°C. Cells were then lysed in either GST lysis buffer (50mM Tris-HCl, pH7.5, 300 mM NaCl, 1.25mM EGTA, 1mM DTT and 0.1% NP-40) or 6xHis-SUMO lysis buffer (50mM Tris-HCl, pH7.5, 300 mM NaCl, 1.25mM EGTA, 1mM DTT, 0.1%NP-40 and 15 mM imidazole) by sonication 6 times for 15 s each. The resultant lysates were then centrifuged at 24,000 X g for 30 minutes at 4°C. The supernatants were then incubated with either Glutathione Sepharose 4B (GE Healthcare, Chicago, IL, USA) or Complete His-Tag Purification Resin (Roche, Basel, Switzerland), that had been prewashed with respective lysis buffer, for 1 hour at 4°C. The beads were then washed five times with respective lysis buffer. GST-tagged proteins were kept with Glutathione Sepharose 4B and resuspended in Bead storage buffer (50mM Tris-HCl, pH7.5, 5 mM MgCl_2_, 25% glycerol), while 6xHis-SUMO-tagged proteins were eluted by elution buffer (50mM Tris-HCl, pH7.5, 300 mM NaCl, 1.25mM EGTA, 1mM DTT, 0.1% NP-40 and 300mM imidazole). Protein concentrations were determined by standard curve intensity measurements from Coomassie blue-stained bovine serum albumin (A7617, Sigma Aldrich, St. Louis, MO, USA) of known concentrations.

For in vitro binding, 3.5 μg of either GST, GST-Elm1^WT^, GST-Elm1^KD^, GST-Elm1_1-420_, GST-Elm1_421-640_, GST-Elm1^S519A^, GST-Elm1^S519D^, GST-Elm1 Elm1^7A^, or GST-Elm1 Elm1^7D^ was incubated with 3.5 μg of either 6xHis-SUMO, 6x-His-SUMO-Gin4, or 6x-His-SUMO-Gin4^KD^ for 1 hour with rotation at in binding buffer (20 mM MOPS, pH7.0, 1 mM EGTA, 100 mM NaCl, 1 mM DTT, 0.1%NP-40). Beads were then washed five times with fresh binding buffer before being extracted with 30 µL of 2X Laemmli Buffer (Bio-Rad Laboratories, Hercules, CA, USA). For Figure 5 E, 5 µL were separated via SDS-PAGE, then the SDS-PAGE gel was cut into two between 100-150kDa, the top half was transferred to PVDF membrane, and blotted with anti-6xHis antibody (1:5000 dilution) followed by HRP-labeled secondary antibody and ECL reagents from the Pierce Fast Western kit (Thermo Scientific, Waltham, MA, USA). The bottom half was stained with SimplyBlue Safe stain (Life Technology, Carlsbad, CA, USA). For Figures 4 B and 4 F, 5μL were separated via SDS-PAGE and stained with SimplyBlue safe stain while 3 μL were separated via SDS-PAGE on a separate gel and transferred to a PVDF membrane before immunoblotting with the anti-6xHis antibody and visualized as described above.

### Yeast protein isolation and immunoblotting

Yeast strains YEF9327, YEF10749, and YEF10750 were inoculated into 25mL of YM-1 media and grown overnight to exponential phase before being pelleted at 4,000 X g for 5 minutes. Cells were then resuspended in 200μL of 4x Laemmli Buffer (Bio-Rad Laboratories), moved to a screw cap tube and lysed via glass beads using a bead beater (BioSpec Mini-Beadbeater 16, Bartlesville, OK, USA) for two rounds of one minute beating and one minute resting on ice. A hole was then poked in the bottom of the tube, the tube was placed into a 15mL conical tube and centrifuged at 3000 X g to collect the lysate. Samples were then diluted 1:4 with water and separated via SDS-PAGE, transferred to a PVDF membrane, before being immunoblotted with mouse anti-GFP (1:3000 dilution, 902601, BioLegend, San Diego, CA, USA) and mouse anti-Cdc28 (1:3000 dilution, sc-515762, Santa Cruz, Dallas, TX, USA) primary antibodies, and HRP-labeled secondary antibody and ECL reagents from the Pierce Fast Western kit (Thermo Scientific).

### Mass spectrometry sample preparation, instrumentation, and analysis

For mass spectrometry of proteins isolated from yeast cultures (our in vivo sample set), yeast strains YEF 9327, YEF10749, and YEF10750 were inoculated into 100mL of YM-1 and allowed to grow to exponential phase overnight at 25°C. Samples were lysed in 200μL RIPA buffer (10mM Tris-Cl pH7.5, 150mM NaCl, 0.5mM EDTA, 0.1% SDS, 1% Triton X100, and 1% deoxycholate), before being diluted with 300μL dilution buffer (10mM Tris-Cl pH7.5, 150mM NaCl, 0.5mM EDTA). These samples were then incubated with 25ul GFP trap beads (ChromoTek GFP-Trap@ Agarose,gta-20, Planegg, Germany) at 4°C for 3 hours, washed with four times in high salt wash buffer (10mM Tris-Cl pH7.5, 300mM NaCl, 0.5mM EDTA, 0.05% NP-40) then 1 time in low salt wash buffer (10mM Tris-Cl pH7.5, 150mM NaCl, 0.5mM EDTA, 0.05% NP-40). Ten percent of the sample was separated via SDS-PAGE, transferred to a PVDF membrane, and immunoblotted with mouse anti-GFP (1:3000 dilution, BioLegend) and HRP-labeled secondary antibody and ECL reagents from the Pierce Fast Western kit (Thermo Scientific), to confirm efficient immunoprecipitation. The remaining sample was separated via SDS-PAGE and stained with SimplyBlue Safe stain (Life Technology). For mass spectrometry of recombinant purified proteins (our in vitro dataset), GST-Elm1^KD^ and 6xHis-SUMO-Gin4 were purified as described above. 1μg of GST-Elm1 was incubated with either buffer (no kinase sample) or 0.2μg 6xHis-SUMO-Gin4 were then incubated in the presence of 2mM ATP for 30 minutes at 30°C with shaking before being separated via SDS-PAGE and stained with SimplyBlue Safe stain.

The gel bands that corresponded to the area overlayed by immunoblot described above (in vivo) or predicted size of GST-Elm1 and 6xHis-SUMO-Gin4 (in vitro) were excised, reduced with Tris (2-carboxyethyl) phosphine hydrochloride (TCEP), alkylated with iodoacetamide, and digested with trypsin. Tryptic digests were analyzed using a standard 90-minute LC gradient on the Thermo Q Exactive Plus (Thermo Fisher, Waltham, MA, USA) mass spectrometer.

Mass spectrometry data were searched with full tryptic specificity against the Swiss-Prot *S. cerevisiae* database (07/26/2021) and a common contaminant database using MaxQuant 1.6.3.3. Variable modifications searched include: Acetylation (+42.01056) on protein N-terminus, Oxidation (+15.99491) on M, Deamidation (+0.98402) on N, and Phosphorylation (+79.9663) on S,T,Y. Proteins and peptides were then filtered by removing common contaminants and incorrect protein modifications. Protein quantification was performed using Razor + unique peptides. Razor peptides are shared (non-unique) peptides assigned to the protein group with the most other peptides (Occam’s razor principle). The abundance of the protein and peptides in a sample was measured from the Intensity (sum of the peptide MS peak areas for the protein). Phosphorylated residues were the primary modifications sought and were identified and filtered out of the dataset and their abundance was compared between the no kinase added sample and in the presence of active Gin4.

### Supplemental material

Four supplemental figures are presented with accompanying figure legends. Figure S1 depicts Elm1 and Gin4 localization in the same strain of yeast. Figure S2 outlines in vivo mass spectrometry results as well as detailed Gin4 auto-phosphorylation sites via in vitro mass spectrometry. Figure S3 depicts localization kinetics and intensity measurements of the Elm1S519A and Elm1S519D mutant proteins tagged with GFP in relation to the septins. Figure S4 depicts the growth patterning and localization of the Elm1 phosphorylation site mutants in the absence of *BNI5*. One supplemental table is provided that shows strains used for the study and a detailed explanation of their construction in the footnotes.

## DATA AVAILABILITY STATEMENT

The data are available from the primary corresponding author (Erfei Bi via) upon request.

## ACKNOWLEDGEMENTS

The authors would like to thank the members of the Bi lab for critical discussions of data interpretation and figure design. We would like to thank former undergraduates Tanya Gupta and Alaina Hunt for help in yeast strain construction and initial preliminary data acquisition. We would also like to thank the Proteomics and Metabolomics Facility at the Wistar Institute for mass spectrometry work and analysis. We would also like to thank J. Hegemann (Heinrich-Heine-Universität Düsseldorf, Germany) and Masayuki Onishi in the lab of John Pringle for generously sharing plasmids. The authors declare no competing financial interests. This work is supported by the National Institutes of Health grants GM116876 (to E.B.)

## AUTHOR CONTRIBUTIONS

J.M. and X.C. conducted the experiments; J.M., X.C., and E.B. designed the experiments; and J.M., X.C., and E.B wrote the manuscript.

## ABBREVIATIONS

GBP: GFP nanobody binding peptide
HDR: hourglass to double ring
IR: inter-region
KA1: kinase associated 1
KD: kinase dead
LC-MS: liquid chromatography mass spectrometry
PAK: p21-activated protein kinase
SAP: septin-associated protein
SC: Synthetic complete minimal media
SUMO: small ubiquitin-like modifier

