## Supplemental Figures S1-S4 and Tables S1 for "Elucidating the Synergistic Role of Elm1 and Gin4 Kinases in Regulating Septin Hourglass Assembly"

**Figure S1. Gin4 localizes to the bud neck prior to Elm1 during bud emergence and maintains localization after Elm1 prior to cytokinesis.**

(A) Montages of representative YEF10802 cells showing maximum-intensity projections of Elm1-GFP in green and Gin4-mScarlet mRuby2-Tub1 in magenta from 12 minutes before to 40 minutes after bud emergence from time-lapse series taken with a 2-minute interval. T = 0 is bud emergence. Scale bars = 1 μm.

(B) Quantification of cells in Figure S1 A. Shown is background subtracted intensity of Elm1-GFP in green and Gin4-mScarlet mRuby2-Tub1 in magenta relative to the maximum value measured from the sum projection of given number cells. The mean is plotted with error bars being the standard deviation.

(C) Montages of representative YEF10802 cells showing maximum-intensity projections of Elm1-GFP in green and Gin4-mScarlet mRuby2-Tub1 in magenta from 16 minutes before to 10 minutes after mitotic spindle break from time-lapse series taken with a 2-minute interval. T = 0 is mitotic spindle break. Scale bars = 1 μm.

(D) Quantification of cells in Figure S1 C. Shown is background subtracted intensity of Elm1-GFP in green and Gin4-mScarlet mRuby2-Tub1 in magenta relative to the maximum value measured from the sum projection of given number cells. The mean is plotted with error bars being the standard deviation.

**Figure S2. Elm1 exhibits Gin4-dependent phosphorylation in vivo and Gin4 heavily autophosphorylates in vitro.**

(A) Analysis of immunoprecipitated Elm1-GFP from control (YEF9327, no GFP), WT (YEF10749, *ELM1-GFP*), and *gin4*Δ (YEF10750, *gin4*Δ *ELM1-GFP*) whole cell lysates separated via SDS-PAGE and immunoblotted with anti-GFP antibody (left) or Coomassie Blue stained (right). Yellow boxes indicate gel area excised for mass spectrometry analysis. IP = immunoprecipitated, IB = immunoblotted.

(B) Protein schematic of Elm1 with indicated domain boundaries labeled with amino acid positions. Green values indicate the position of a phosphorylated residue discovered via mass spectrometry only in the WT Elm1-GFP IP sample. Red values indicate the position of a phosphorylated residue found enriched in the *gin4*Δ Elm1-GFP IP mass spectrometry sample compared to that of the WT sample. Blue values indicate the position of a phosphorylated residue found enriched in the WT Elm1-GFP IP mass spectrometry sample compared to that of the *gin4*Δ sample.

(C) Phosphorylated residues discovered in the 6xHis-SUMO-Gin4 protein after kinase reaction in the presence of GST-Elm1$^{KD}$. Asterisk (*) indicates phosphorylated residues believed to be Elm-dependent from a previous study (Asano et al., 2006).

**Figure S3. Phosphorylation at S519 in Elm1 is not necessary for Elm1 localization to the bud neck.**

(A) Montages of representative cells of Elm1$^{WT}$, Elm1$^{S519A}$, and Elm1$^{S519D}$ tagged with GFP in green and Cdc3-mCherry shown in magenta. The images show maximum-intensity projections of the indicated fluorescent protein from 20 minutes before to 40 minutes after bud emergence with selected frames from time-lapse series taken with a 2-min interval. Strains used are as follows: YEF11688 (*GFP-ELM1$^{WT}$ CDC3-mCherry*), YEF11689 (*GFP-elm1$^{S519A}$ CDC3-mCherry*), and YEF11690 (*GFP-elm1$^{S519D}$ CDC3-mCherry*). Scale bars = 2 μm.

(B) Quantification of the cells in Figure S3 A. Top panel shown is background subtracted integrated intensity measured of GFP-Elm1 from the sum projection of a given number of cells for each strain. The mean is plotted with error bars being standard deviation. A.U. = arbitrary units. Bottom panel shown is background subtracted integrated intensity measured of Cdc3-mCherry (open circles) and GFP-Elm1 (crosses) in given number of cells per strain at 20 minutes after bud emergence. Each plotted point is a single cell's bud neck measured intensity. N.S. = not significant ($p > 0.05$) by unpaired Student's t-test.

(C) Montages of representative cells of Elm1$^{WT}$, Elm1$^{S519A}$, and Elm1$^{S519D}$ tagged with GFP in green, and Cdc3-mCherry shown in magenta. The images show maximum-intensity projections of the indicated fluorescent protein from 40 minutes before to 8 minutes after septin hourglass to double ring transition with selected frames from time-lapse series taken with a 2-min interval. Strains used are the same as listed in Figure S3 A. Scale bars = 2 μm.

(D) Quantification of the cells in Figure S3 C. Top panel shown is background subtracted integrated intensity measured of GFP-Elm1 from the sum projection of a given number of cells for each strain. The mean is plotted with error bars being standard deviation. A.U. = arbitrary units. Bottom panel shown is background subtracted integrated intensity measured of Cdc3-mCherry (open circles) and GFP-Elm1 (crosses) in given number of cells per strain at 20 minutes after bud emergence. Each plotted point is a single cell's bud neck measured intensity. N.S. = not significant ($p > 0.05$), single asterisk = $p < 0.05$, and double asterisk = $p < 0.01$ by unpaired Student's t-test

(E) Western Blot analysis of Elm1 phospho-site mutant protein expression compared to Elm1$^{WT}$. Strains used are as follows: YEF11665 (*GFP-ELM1*$^{WT}$), YEF11666 (*GFP-elm1*$^{S519A}$), YEF11667 (*GFP-elm1*$^{S519D}$), YEF11668 (*GFP-elm1*$^{7A}$) YEF11669 (*GFP-elm1*$^{S519D}$), and YEF2232 (no GFP control). Top: immunoblotted with antibody anti-GFP; Bottom: immunoblotted with antibody anti-Cdc28 as the loading control. This experiment was repeated 3 times.

**Figure S4. Deletion of *BNI5* does not exacerbate the cell growth and morphologies of Elm1 phosphorylation mutants.**

(A) The sensitivity of *bni5*Δ cells with *elm1* mutants was tested on synthetic complete (SC) plates. Ten-fold serial dilutions of YEF11761 (*bni5*Δ *GFP-ELM1*$^{WT}$ *CDC3-mCherry*), YEF11762 (*bni5*Δ *GFP-elm1*$^{S519A}$ *CDC3-mCherry*), YEF11763 (*bni5*Δ *GFP-elm1*$^{S519D}$ *CDC3-mCherry*), YEF11764 (*bni5*Δ *GFP-elm1*$^{7A}$ *CDC3-mCherry*), and YEF11765 (*bni5*Δ *GFP-elm1*$^{7D}$ *CDC3-mCherry*) cultures were spotted on SC plates and incubated at 25ºC (left) and for 37ºC (right) 3 days.

(B and C) Representative images of the indicated *bni5*Δ cells were cultured at 25ºC (B) and 37ºC (C) overnight with bright field (top) and maximum intensity projections of GFP-tagged Elm1 mutants shown in green (middle) and Cdc3-mCherry shown in magenta (bottom). Strains used are listed in Figure S4 A. The dashed line indicates the cell periphery. Scale bars = 2 μm.

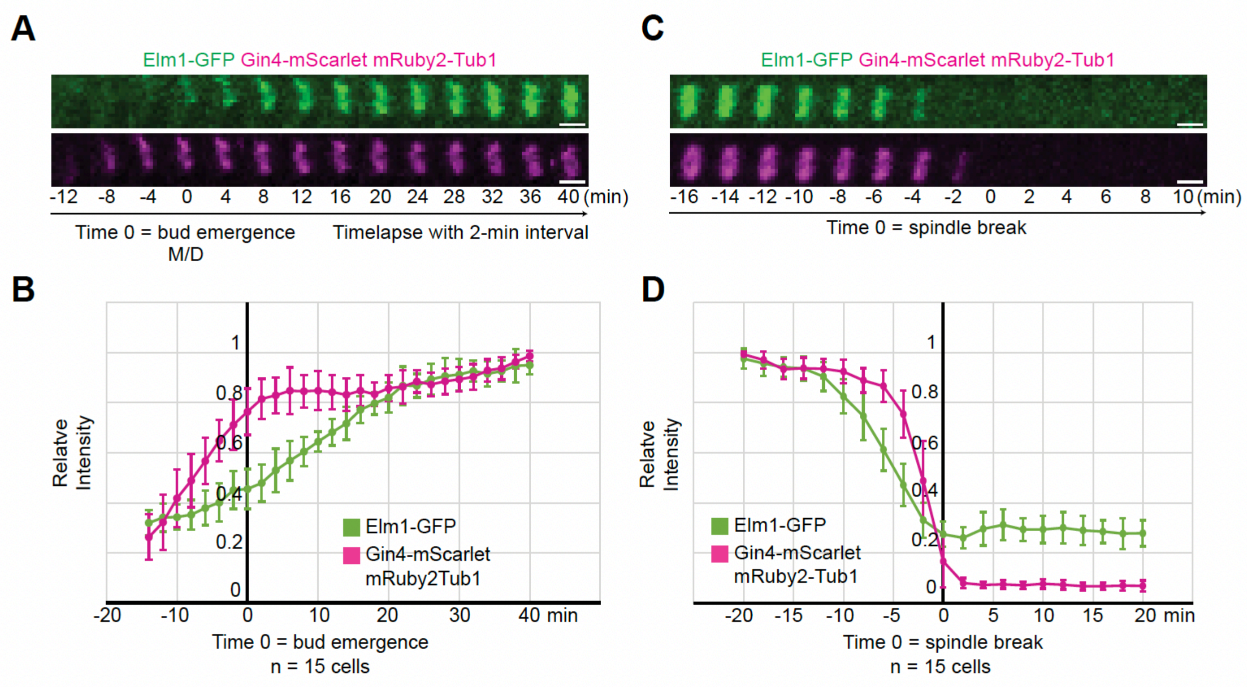


Figure S1. Marquardt et al.

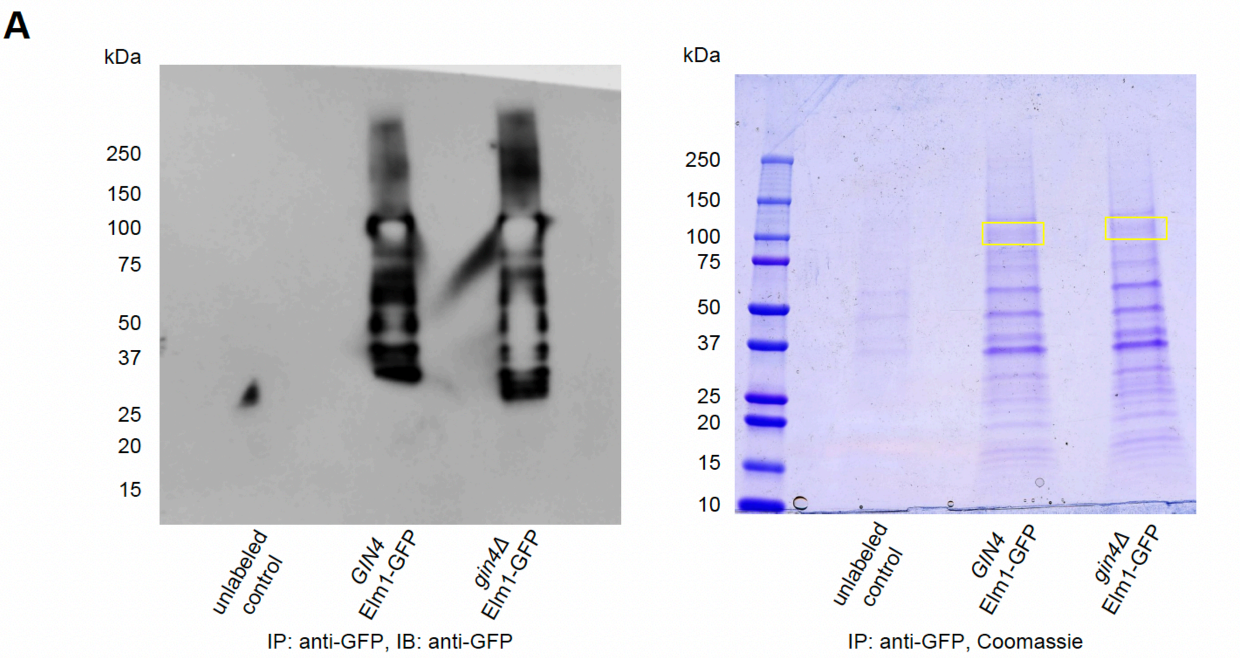


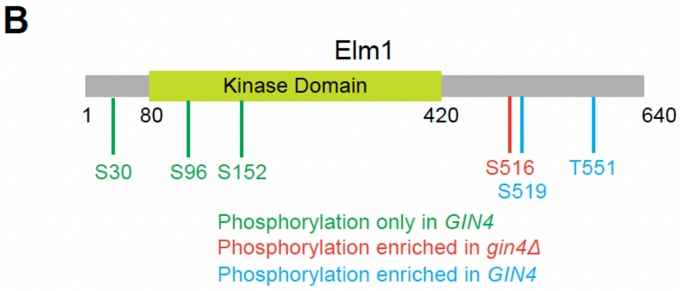


**C** In vitro Phosphorylation Sites in His-SUMO-Gin4

| | | |
|---|---|---|
| S40 | S439 | S666 |
| T63 | S448 | T669 |
| S64 | S453* | S673 |
| S68 | T454 | T685 |
| T69 | S459 | S714 |
| T309 | S460* | S719 |
| S382 | T462* | S720 |
| S384 | S471 | S736 |
| S385 | S483 | S801 |
| S387 | S486 | S807 |
| S389* | T487 | S812 |
| S391 | T488 | S818 |
| S406 | S502 | T819 |
| S409 | S506 | S822 |
| T411 | S518 | T824 |
| S413 | T519 | S842 |
| S419 | T539* | S889 |
| S426 | S540* | S947 |
| S429 | S541 | S949 |
| T435 | S617* | S950 |
| S438 | S639* | S1029 |

Figure S2. Marquardt et al.

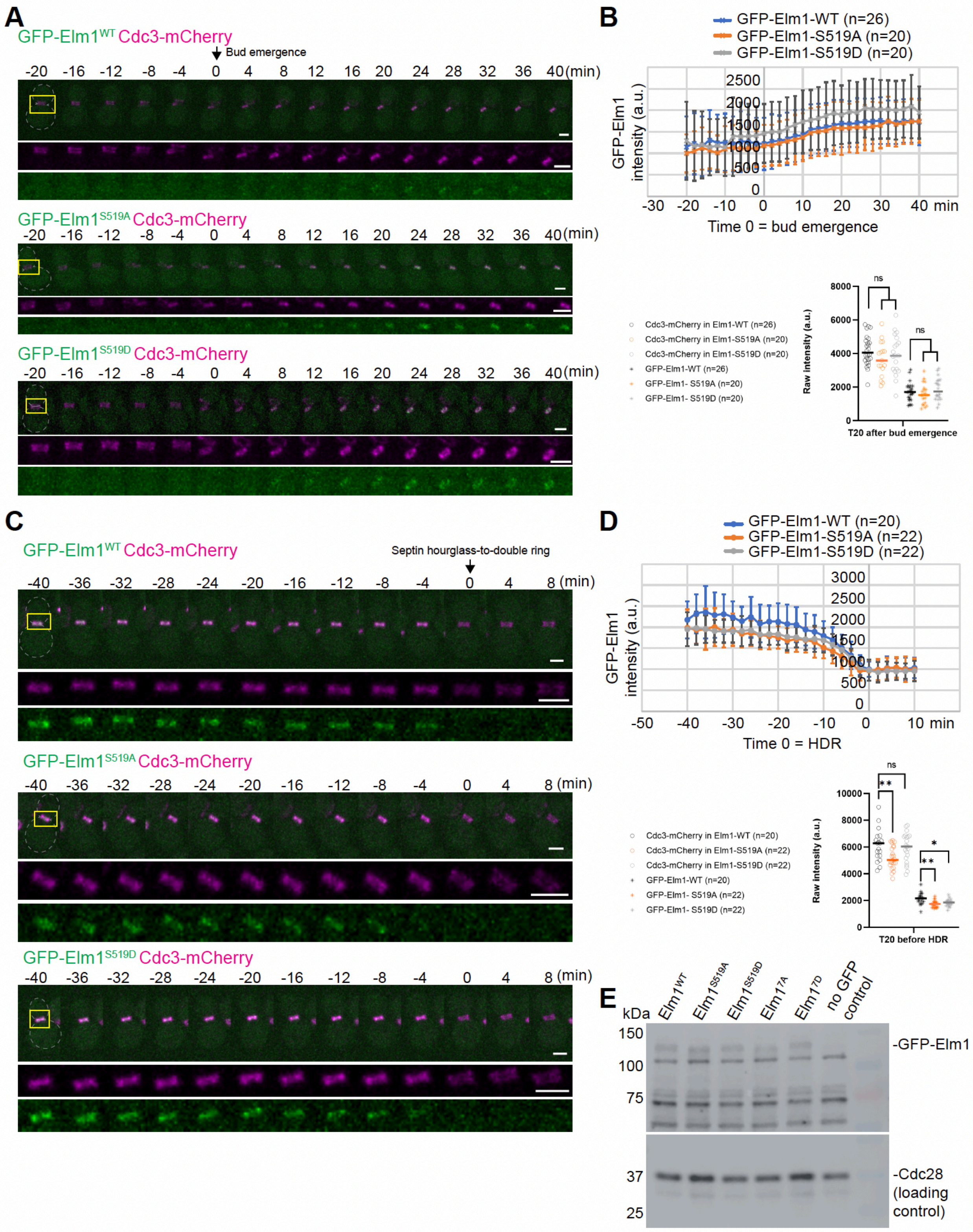


Figure S3. Marquardt et al.

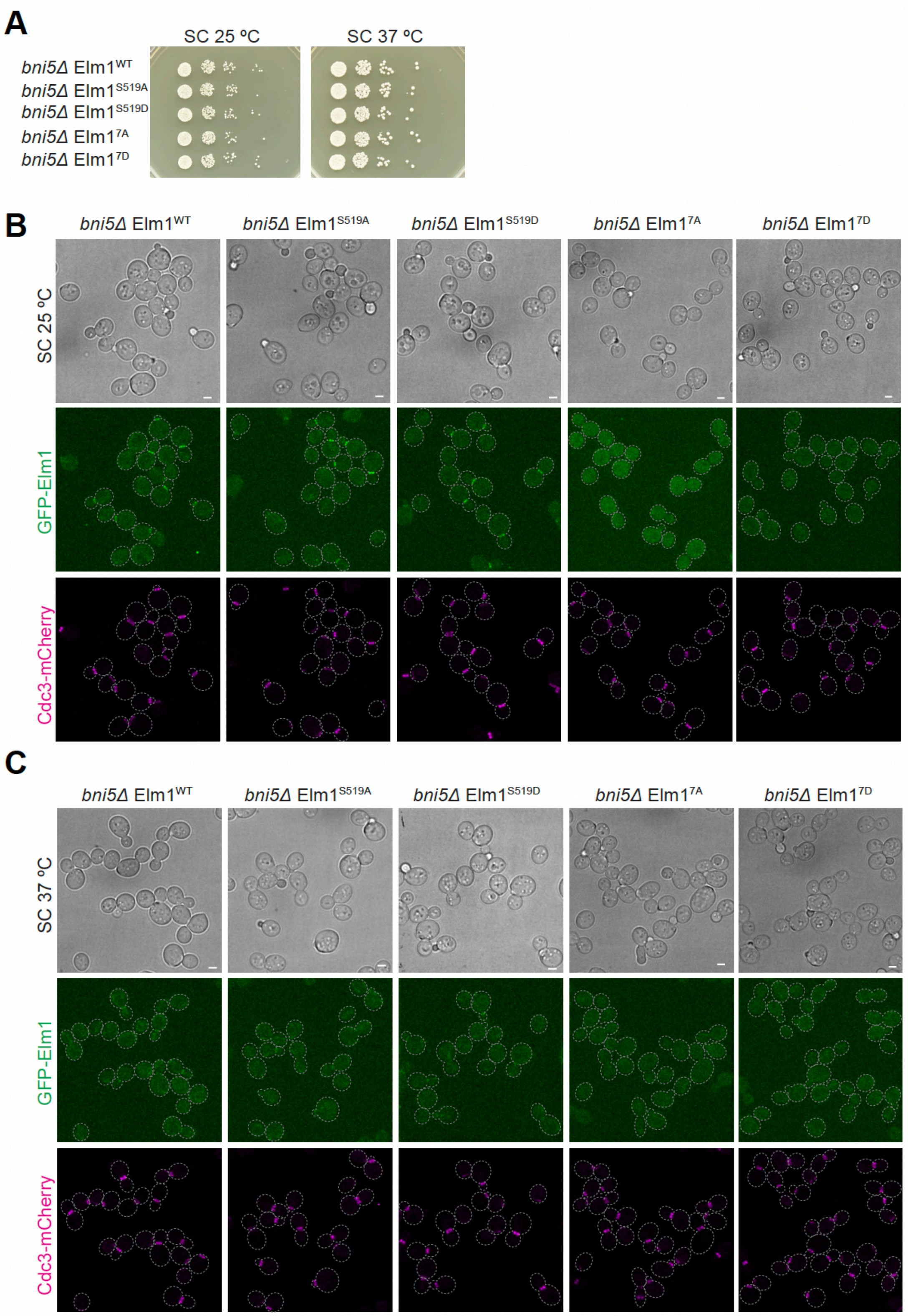


Figure S4. Marquardt et al.

| Strain | Genotype | Source |
|---|---|---|
| YEF473A | **a** *his3 leu2 lys2 trp1 ura3* | (Bi and Pringle, 1996) |
| YEF8438 | As YEF473A except *shs1Δ::TRP1 CDC3-GFP-LEU2 pHis3-mRuby2-TUB1-URA3* | This study[a] |
| YEF9180 | As YEF473A except *CDC3-GFP-LEU2 pHis3-mRuby2-TUB1-HPH* | (Marquardt et al., 2020) |
| YEF9305 | As YEF473A except *ELM1-GFP-CaURA3 CDC3-mCherry-LEU2* | This study[b] |
| YEF9327 | As YEF473A except *bar1Δ::HIS3MX6* | (Marquardt et al., 2020) |
| YEF9335 | As YEF473A except *elm1*$^{KD(K117R)}$*-mApple-GBP-CaURA3* | (Marquardt et al., 2020) |
| YEF9362 | As YEF473A except *SHS1-GFP-HIS3 elm1*$^{KD(K117R)}$*-mApple-GBP-CaURA3* | (Marquardt et al., 2020) |
| YEF9448 | As YEF473A except *ELM1-mApple-GBP-CaURA3* | This study[c] |
| YEF9485 | As YEF473A except *gin4Δ::TRP1 ELM1-mApple-GBP-CaURA3* | This study[d] |
| YEF9641 | As YEF473A except *gin4Δ::TRP1 CDC3-GFP-LEU2 pHis3-mRuby2-TUB1-HPH* | This study[d] |
| YEF9935 | As YEF473A except *elm1Δ::KanMX6 CDC3-GFP-LEU2 pHis3-mRuby2-TUB1-HPH* | This study[e] |
| YEF10238 | As YEF473A except *elm1Δ::KanMX6 gin4Δ::TRP1 CDC3-GFP-LEU2 pHis3-mRuby2-TUB1-HPH* | This study[e] |
| YEF10440 | As YEF473A except *ELM1-GFP-SpHIS5 CDC3-mCherry-LEU2* | This study[f] |
| YEF10441 | As YEF473A except *GIN4-GFP-SpHIS5 CDC3-mCherry-LEU2* | This study[g] |
| YEF10460 | As YEF473A except *gin4Δ::TRP1 ELM1-GFP-SpHIS5 CDC3-mCherry-LEU2* | This study[d] |
| YEF10558 | As YEF473A except *GIN4-GFP-SpHIS5 CDC3-mCherry-LEU2 pHis3-mRuby2-TUB1-HPH* | This study[h] |
| YEF10559 | As YEF473A except *elm1Δ::KanMX6 GIN4-GFP-SpHIS5 CDC3-mCherry-LEU2 pHis3-mRuby2-TUB1-HPH* | This study[i] |
| YEF10665 | As YEF473A except *SHS1-GFP-HIS3 ELM1-mApple-GBP-CaURA3* | This study[j] |
| YEF10666 | As YEF473A except *gin4Δ::TRP1 SHS1-GFP-HIS3 ELM1-mApple-GBP-CaURA3* | This study[j] |

| | | |
|---|---|---|
| YEF10672 | As YEF473A except *gin4*$^{\Delta KA1}$ *ELM1-GFP-SpHIS5 CDC3-mCherry-LEU2* | This study[k] |
| YEF10673 | As YEF473A except *ELM1-GFP-SpHIS5 gin4*$^{KD(K48A)}$*-mScarlet-KanMX6* | This study[l] |
| YEF10738 | As YEF473A except *gin4Δ::TRP1 elm1*$^{KD(K117R)}$*-mApple-GBP-CaURA3* | This study[d] |
| YEF10743 | As YEF473A except *gin4Δ::TRP1 SHS1-GFP-HIS3 elm1*$^{KD(K117R)}$*-mApple-GBP-CaURA3* | This study[d] |
| YEF10749 | As YEF473A except *bar1Δ::HIS3MX6 ELM1-GFP-CaURA3* | This study[m] |
| YEF10750 | As YEF473A except *gin4Δ::TRP1 bar1Δ::HIS3MX6 ELM1-GFP-CaURA3* | This study[n] |
| YEF10802 | As YEF473A except *ELM1-GFP-SpHIS5 GIN4-mScarlet-CaURA3 pHis3-mRuby2-TUB1-HPH* | This study[o] |
| YEF10989 | As YEF473A except *elm1Δ::KanMX6 gin4*$^{KD(K48A)}$*-GFP-SpHIS5 CDC3-mCherry-LEU2* | This study[p] |
| YEF10990 | As YEF473A except *elm1Δ::KanMX6 gin4*$^{\Delta KA1}$ *CDC3-GFP-LEU2 pHis3-mRuby2-TUB1-HPH* | This study[q] |
| YEF11144 | As YEF473A except *elm1Δ::KanMX6 gin4*$^{KD(K48A)}$ *CDC3-GFP-LEU2 pHis3-mRuby2-TUB1-HPH* | This study[r] |
| YEF11454 | As YEF473A except *shs1Δ::TRP1 GIN4-GFP-SpHIS5 CDC3-mCherry-LEU2 pHis3-mRuby2-TUB1-HPH* | This study[a] |
| YEF11493 | As YEF473A except *SHS1-GFP-SpHIS5 GIN4-mScarlet-CaURA3 pHis3-mRuby2-TUB1-HPH* | This study[s] |
| YEF11494 | As YEF473A except *elm1*$^{KD(K117R)}$*-GBP-KanMX6 SHS1-GFP-SpHIS5 GIN4-mScarlet-CaURA3 pHis3-mRuby2-TUB1-HPH* | This study[s] |
| YEF11495 | As YEF473A except *ELM1-GBP-KanMX6 SHS1-GFP-SpHIS5 GIN4-mScarlet-CaURA3 pHis3-mRuby2-TUB1-HPH* | This study[t] |
| YEF11496 | As YEF473A except *elm1*$^{KD(K117R)}$ *SHS1-GFP-SpHIS5 GIN4-mScarlet-CaURA3 pHis3-mRuby2-TUB1-HPH* | This study[u] |
| YEF11606 | As YEF473A except *elm1*$_{1-420}^{KD(K117R)}$*-GBP-KanMX6 SHS1-GFP-SpHIS5 GIN4-mScarlet-CaURA3 pHis3-mRuby2-TUB1-HPH* | This study[s] |
| YEF11607 | As YEF473A except *elm1*$_{1-420}$*-GBP-KanMX6 SHS1-GFP-SpHIS5 GIN4-mScarlet-CaURA3 pHis3-mRuby2-TUB1-HPH* | This study[s] |
| YEF11665 | As YEF473A except *GFP-ELM1* | This study[v] |
| YEF11666 | As YEF473A except *GFP-elm1*$^{S519A}$ | This study[v] |

| | | |
|---|---|---|
| YEF11667 | As YEF473A except *GFP-elm1$^{S519D}$* | This study[v] |
| YEF11668 | As YEF473A except *GFP-elm1$^{7A}$* | This study[v] |
| YEF11669 | As YEF473A except *GFP-elm1$^{7D}$* | This study[v] |
| YEF11679 | As YEF473A except *GFP-ELM1 pHis3-mRuby2-TUB1-HPH* | This study[h] |
| YEF11680 | As YEF473A except *GFP-elm1$^{S519A}$ pHis3-mRuby2-TUB1-HPH* | This study[h] |
| YEF11681 | As YEF473A except *GFP-elm1$^{S519D}$ pHis3-mRuby2-TUB1-HPH* | This study[h] |
| YEF11682 | As YEF473A except *GFP-elm1$^{7A}$ pHis3-mRuby2-TUB1-HPH* | This study[h] |
| YEF11683 | As YEF473A except *GFP-elm1$^{7D}$ pHis3-mRuby2-TUB1-HPH* | This study[h] |
| YEF11688 | As YEF473A except *GFP-ELM1 CDC3-mCherry-LEU2* | This study[w] |
| YEF11689 | As YEF473A except *GFP-elm1$^{S519A}$ CDC3-mCherry-LEU2* | This study[w] |
| YEF11690 | As YEF473A except *GFP-elm1$^{S519D}$ CDC3-mCherry-LEU2* | This study[w] |
| YEF11691 | As YEF473A except *GFP-elm1$^{7A}$ CDC3-mCherry-LEU2* | This study[w] |
| YEF11692 | As YEF473A except *GFP-elm1$^{7D}$ CDC3-mCherry-LEU2* | This study[w] |
| YEF11708 | As YEF473A except *gin4Δ::TRP1 GFP-ELM1 CDC3-mCherry-LEU2* | This study[d] |
| YEF11709 | As YEF473A except *gin4Δ::TRP1 GFP-elm1$^{S519A}$ CDC3-mCherry-LEU2* | This study[d] |
| YEF11710 | As YEF473A except *gin4Δ::TRP1 GFP-elm1$^{S519D}$ CDC3-mCherry-LEU2* | This study[d] |
| YEF11711 | As YEF473A except *gin4Δ::TRP1 GFP-elm1$^{7A}$ CDC3-mCherry-LEU2* | This study[d] |
| YEF11712 | As YEF473A except *gin4Δ::TRP1 GFP-elm1$^{7D}$ CDC3-mCherry-LEU2* | This study[d] |
| YEF11761 | As YEF473A except *bni5Δ::HIS3MX6 GFP-ELM1 CDC3-mCherry-LEU2* | This study[x] |
| YEF11762 | As YEF473A except *bni5Δ::HIS3MX6 GFP-elm1$^{S519A}$ CDC3-mCherry-LEU2* | This study[x] |
| YEF11763 | As YEF473A except *bni5Δ::HIS3MX6 GFP-elm1$^{S519D}$ CDC3-mCherry-LEU2* | This study[x] |
| YEF11764 | As YEF473A except *bni5Δ::HIS3MX6 GFP-elm1$^{7A}$ CDC3-mCherry-LEU2* | This study[x] |
| YEF11765 | As YEF473A except *bni5Δ::HIS3MX6 GFP-elm1$^{7D}$ CDC3-mCherry-LEU2* | This study[x] |
| YEF11766 | As YEF473A except *shs1Δ::TRP1 GFP-ELM1 CDC3-mCherry-LEU2* | This study[a] |

| | | |
|---|---|---|
| YEF11767 | As YEF473A except *shs1Δ::TRP1 GFP-elm1$^{S519A}$ CDC3-mCherry-LEU2* | This study[a] |
| YEF11768 | As YEF473A except *shs1Δ::TRP1 GFP-elm1$^{S519D}$ CDC3-mCherry-LEU2* | This study[a] |
| YEF11769 | As YEF473A except *shs1Δ::TRP1 GFP-elm1$^{7A}$ CDC3-mCherry-LEU2* | This study[a] |
| YEF11770 | As YEF473A except *shs1Δ::TRP1 GFP-elm1$^{7D}$ CDC3-mCherry-LEU2* | This study[a] |

**Table S1. Strains used in this study.**

[a] A DNA fragment carrying *shs1Δ::TRP1* was amplified by PCR using the chromosomal DNA from YEF7506 (our lab stock) as the template DNA and the pair of primers Shs1-F-check and Shs1-R-check, and then transformed into YEF8102 (our lab stock), YEF10558, YEF11688, YEF11689, YEF11690, YEF11691, and YEF11692 to generate YEF8438, YEF11454, YEF11766, YEF11767, YEF11768, YEF11769, and YEF11770, respectively.

[b] BglII-digested YIp128-CDC3-mCherry (integrated at the *CDC3* locus) (Gao et al., 2007) was transformed into YEF473A to generate YEF5804. Then a DNA fragment carrying *ELM1-GFP-CaURA3* was amplified by PCR using the chromosomal DNA from YEF8299 (Marquardt et al., 2020) as the template DNA and the pair of primers Elm1-Ftag-check and Elm1-R-check, and then transformed into YEF5804 to generate YEF9305.

[c] A DNA fragment carrying *ELM1-mApple-GBP-CaURA3* was amplified by PCR using the chromosomal DNA from YEF9335 (Marquardt et al., 2020) as the template DNA and the pair of primers Elm1-Ftag-check and Elm1-R-check, and then transformed into YEF473A to generate YEF9448.

[d] A DNA fragment carrying *gin4Δ::TRP1* was amplified by PCR using the chromosomal DNA from YEF7267 (our lab stock) as the template DNA and the pair of primers Gin4-F-check and Gin4-R-check, and then transformed into YEF9448, YEF9180, YEF10440, YEF9335, YEF9362, YEF11688, YEF11689, YEF11690, YEF11691, and YEF11692 to generate YEF9485, YEF9641, YEF10460, YEF10738, YEF10743, YEF11708, YEF11709, YEF11710, YEF11711, and YEF11712, respectively.

[e] A DNA fragment carrying *elm1Δ::KanMX6* was amplified by PCR using the chromosomal DNA from YEF7515 (our lab stock) as the template DNA and the pair of primers Elm1-F-check and Elm1-R-check, and then transformed into YEF9180 and YEF9641 to generate YEF9935 and YEF10238, respectively.

[f] A DNA fragment carrying *ELM1-GFP-SpHIS5* was amplified by PCR using the chromosomal DNA from YEF8250 (Marquardt et al., 2020) as the template DNA and the pair of primers Elm1-Ftag-check and Elm1-R-check, and then transformed into YEF5804 to generate YEF10440.

[g] A DNA fragment carrying *GIN4-GFP-SpHIS5* was amplified by PCR using the chromosomal DNA from YEF8249 (our lab stock) as the template DNA and the pair of primers Gin4-Ftag-check and Gin4-R-check, and then transformed into YEF5804 to generate YEF10441.

[h] BsaBI-digested plasmid pHis3:mRuby2-Tub1+3'UTR::HPH (Markus et al., 2015) was transformed into YEF10441, YEF11665, YEF11666, YEF11667, YEF11668, and YEF11669 to generate strains YEF10558, YEF11679, YEF11680, YEF11681, YEF11682, and YEF11683, respectively.

[i] A DNA fragment carrying *elm1Δ::KanMX6* was amplified by PCR using the chromosomal DNA from YEF7515 (our lab stock) as the template DNA and the pair of primers Elm1-F-check and Elm1-R-check, and then transformed into YEF10441 to generate YEF10476. Then BsaBI-digested plasmid pHis3:mRuby2-Tub1+3'UTR::HPH (Markus et al., 2015) was transformed into YEF10476 to generate strain YEF10559.

[j] A DNA fragment carrying *SHS1-GFP-HIS3* was amplified by PCR using the chromosomal DNA from YEF6651 (our lab stock) as the template DNA and the pair of primers Shs1-Ftag-check and Shs1-R-check, and then transformed into YEF9448 and YEF9485 to generate YEF10665 and YEF10666, respectively.

[k] To generate YEF10672, *GIN4* was first deleted by replacing with URA3-KanMX6 by amplifying *gin4Δ::URA3-KanMX6* by PCR using the plasmid pFA6a-URA3-KanMX6 (a gift from John Pringle) as the template DNA and the pair of primers Gin4-F1 and Gin4-R1, and then transformed into YEF473A to generate YEF8122. Then a DNA fragment carrying *gin4*$^{\Delta KA1}$ was amplified by PCR using the chromosomal DNA from YEF473A as the template DNA and the pair of primers Gin4-F-check and Gin4-dKA1-R, and then transformed into YEF8122 and selected for the loss of the URA3-KanMX6 cassette to generate YEF10585. Then a DNA fragment carrying *ELM1-GFP-SpHIS5* was amplified by PCR using the chromosomal DNA from YEF8250 (Marquardt et al., 2020) as the template DNA and the pair of primers Elm1-Ftag-check and Elm1-R-check, and then transformed into YEF10585 to generate YEF10639. Lastly, BsaBI-digested plasmid pHis3:mRuby2-Tub1+3'UTR::HPH (Markus et al., 2015) was transformed into YEF10639 to generate strain YEF10672.

[l] To generate YEF10673, *GIN4* was first deleted by replacing with URA3-KanMX6 by amplifying *gin4Δ::URA3-KanMX6* by PCR using the plasmid pFA6a-URA3-KanMX6 (a gift from John Pringle) as the template DNA and the pair of primers Gin4-F1 and Gin4-R1, and then transformed into YEF473A to generate YEF8122. Then to generate endogenous *GIN4* with the kinase dead mutation K48A, two DNA fragments were generated by PCR using the chromosomal DNA from YEF473A as the template DNA: fragment 1 used Gin4-F-check and Gin4-K48A-R, fragment 2 used Gin-K48A-F and Gin4-R-check. These two fragments have 20 bases of homology between them via the K48A primers to site directed mutagenize the K amino acid to A. A second amplification by PCR was done using the above two fragments as the template DNA (after gel extraction and purification) and the pair of primers Gin4-F-check and Gin4-R-check to combine the two fragments. This long fragment was then transformed into YEF8122 and selected for the loss of the *URA3-KanMX6* cassette to generate YEF10509. Then a DNA fragment containing *GIN4-mScarlet-KanMX6* was amplified by PCR using the genomic DNA from YEF10065 (our lab stock) as the template DNA and the pair of primers Gin4-Ftag-check and Gin4-R-check, and then transformed into YEF10509 to generate YEF10590. Lastly, a DNA fragment carrying *ELM1-GFP-SpHIS5* was amplified by PCR using the chromosomal DNA from YEF8250 (Marquardt et al., 2020) as the template DNA and the pair of primers Elm1-Ftag-check and Elm1-R-check, and then transformed into YEF10590 to generate YEF10673.

[m] A DNA fragment carrying *ELM1-GFP-CaURA3* was amplified by PCR using the chromosomal DNA from YEF8299 (Marquardt et al., 2020) as the template DNA and the pair of primers Elm1-Ftag-check and Elm1-R-check, and then transformed into YEF9327 to generate YEF10749.

[n] A DNA fragment carrying *gin4Δ::TRP1* was amplified by PCR using the chromosomal DNA from YEF7267 (our lab stock) as the template DNA and the pair of primers Gin4-F-check and Gin4-R-check, and then transformed into YEF9327 to generate YEF10675. Then a DNA fragment carrying *ELM1-GFP-CaURA3* was amplified by PCR using the chromosomal DNA from YEF8299 (Marquardt et al., 2020) as the template DNA and the pair of primers Elm1-Ftag-check and Elm1-R-check, and then transformed into YEF10675 to generate YEF10750.

[o] A DNA fragment carrying *GIN4-mScarlet-I-CaURA3* was amplified by PCR using pFA6a-link-ymScarlet-I-CaURA (Marquardt et al., 2020) as the template and the pair of primers Gin4-F5 and Gin4-R3, and then transformed into YEF473A to generate YEF10757. Then a DNA fragment carrying *ELM1-GFP-SpHIS5* was amplified by PCR using the chromosomal DNA from YEF8250 (Marquardt et al., 2020) as the template DNA and the pair of primers Elm1-Ftag-check and Elm1-R-check, and then transformed into YEF10757 to generate YEF10769. Lastly, BsaBI-digested plasmid pHis3:mRuby2-Tub1+3'UTR::HPH (Markus et al., 2015) was transformed into YEF10769 to generate strain YEF10802.

[p] A DNA fragment carrying *GIN4-GFP-SpHIS5* was amplified by PCR using the chromosomal DNA from YEF8249 (our lab stock) as the template DNA and the pair of primers Gin4-Ftag-check and Gin4-R-check, and then transformed into YEF10509 (see [k] above) to generate YEF10589. Then BglII-digested YIp128-CDC3-mCherry (integrated at the *CDC3* locus) (Gao et al., 2007) was transformed into YEF10589 to generate YEF10646. Lastly, a DNA fragment carrying *elm1Δ::KanMX6* was amplified by PCR using the chromosomal DNA from YEF7515 (our lab stock) as the template DNA and the pair of primers Elm1-F-check and Elm1-R-check, and then transformed into YEF10646 to generate YEF10989.

[q] BglII-digested YIp128-CDC3-GFP (integrated at the *CDC3* locus) (Caviston et al., 2003) was transformed into YEF10585 (see [j] above) to generate YEF10644. Then BsaBI-digested plasmid pHis3:mRuby2-Tub1+3'UTR::HPH (Markus et al., 2015) was transformed into YEF10644 to generate strain YEF10678. Lastly, a DNA fragment carrying *elm1Δ::KanMX6* was amplified by PCR using the chromosomal DNA from YEF7515 (our lab stock) as the template DNA and the pair of primers Elm1-F-check and Elm1-R-check, and then transformed into YEF10678 to generate YEF10990.

[r] BglII-digested YIp128-CDC3-GFP (integrated at the *CDC3* locus) (Caviston et al., 2003) was transformed into YEF10509 (see [k] above) to generate YEF10591. Then BsaBI-digested plasmid pHis3:mRuby2-Tub1+3'UTR::HPH (Markus et al., 2015) was transformed into YEF10591 to generate strain YEF11089. Lastly, a DNA fragment carrying *elm1Δ::KanMX6* was amplified by PCR using the chromosomal DNA from YEF7515 (our lab stock) as the template DNA and the pair of primers Elm1-F-check and Elm1-R-check, and then transformed into YEF11089 to generate YEF11144.

[s] A DNA fragment carrying *GIN4-mScarlet-I-CaURA3* was amplified by PCR using the chromosomal DNA from YEF10757 (see [n] above) as the template and the pair of primers Gin4-Ftag-check and Gin4-R-check, and then transformed into YEF8101 (our lab stock), YEF9855 (Marquardt et al., 2020), YEF9978 (Marquardt et al., 2020), and YEF10000 (Marquardt et al.,

2020) to generate YEF11381, YEF11382, YEF11603, and YEF11604, respectively. Then BsaBI-digested plasmid pHis3:mRuby2-Tub1+3'UTR::HPH (Markus et al., 2015) was transformed into YEF11381, YEF11382, YEF11603, and YEF11604 to generate strain YEF11493, YEF11494, YEF11606, and YEF11607, respectively.

[t] A DNA fragment carrying *ELM1-GBP-KanMX6* was amplified by PCR using the chromosomal DNA from YEF9809 (Marquardt et al., 2020) as the template and the pair of primers Elm1-Ftag-check and Elm1-R-check, and then transformed into YEF8101 (our lab stock) to generate YEF11371. Then a DNA fragment carrying *GIN4-mScarlet-I-CaURA3* was amplified by PCR using the chromosomal DNA from YEF10757 (see [n] above) as the template and the pair of primers Gin4-Ftag-check and Gin4-R-check, and then transformed into YEF11371 to generate YEF11455. Lastly, BsaBI-digested plasmid pHis3:mRuby2-Tub1+3'UTR::HPH (Markus et al., 2015) was transformed into YEF11455 to generate YEF11495.

[u] A DNA fragment carrying *SHS1-GFP-HIS3* was amplified by PCR using the chromosomal DNA from YEF6651 (our lab stock) as the template DNA and the pair of primers Shs1-Ftag-check and Shs1-R-check, and then transformed into YEF9206 (Marquardt et al., 2020) to generate YEF11374. Then a DNA fragment carrying *GIN4-mScarlet-I-CaURA3* was amplified by PCR using the chromosomal DNA from YEF10757 (see [n] above) as the template and the pair of primers Gin4-Ftag-check and Gin4-R-check, and then transformed into YEF11374 to generate YEF11456. Lastly, BsaBI-digested plasmid pHis3:mRuby2-Tub1+3'UTR::HPH (Markus et al., 2015) was transformed into YEF11456 to generate YEF11496.

[v] A DNA fragment containing either *GFP-ELM1*, *GFP-elm1*$^{S519A}$, *GFP-elm1*$^{S519D}$, *GFP-elm1*$^{7A}$, or *GFP-elm1*$^{7D}$ was PCR amplified using pUG36-Elm1, pUG36-Elm1$^{S519A}$, pUG36-Elm1$^{S519D}$, pUG36-Elm1$^{7A}$, or pUG36-Elm1$^{7D}$ (all generated from this study), respectively as the template DNA and the pair of primers F-Elm1-prom and R-Elm1-term-Elm1 (640), and then transformed into YEF8123 (Marquardt et al., 2020) to generate YEF11665, YEF11666, YEF11667, YEF11668, and YEF11669, respectively.

[w] BglII-digested YIp128-CDC3-mCherry (integrated at the *CDC3* locus) (Gao et al., 2007) was transformed into YEF11665, YEF11666, YEF11667, YEF11668, and YEF11669 to generate YEF11688, YEF11689, YEF11690, YEF11691, and YEF11692, respectively.

[x] A DNA fragment containing *bni5Δ::HIS3MX6* was amplified by PCR using the chromosomal DNA from YEF6316 (our lab stock) as the template DNA and the pair of primers Bni5-244bp-US-Start and Bni5-381bp-DS-Stop, and then transformed into YEF11688, YEF11689, YEF11690, YEF11691, and YEF11692 to generate YEF11761, YEF11762, YEF11763, YEF11764, and YEF11765, respectively.
